## Supplementary Figures for "The origin, evolution and molecular diversity of the chemokine system"

Fig. S1

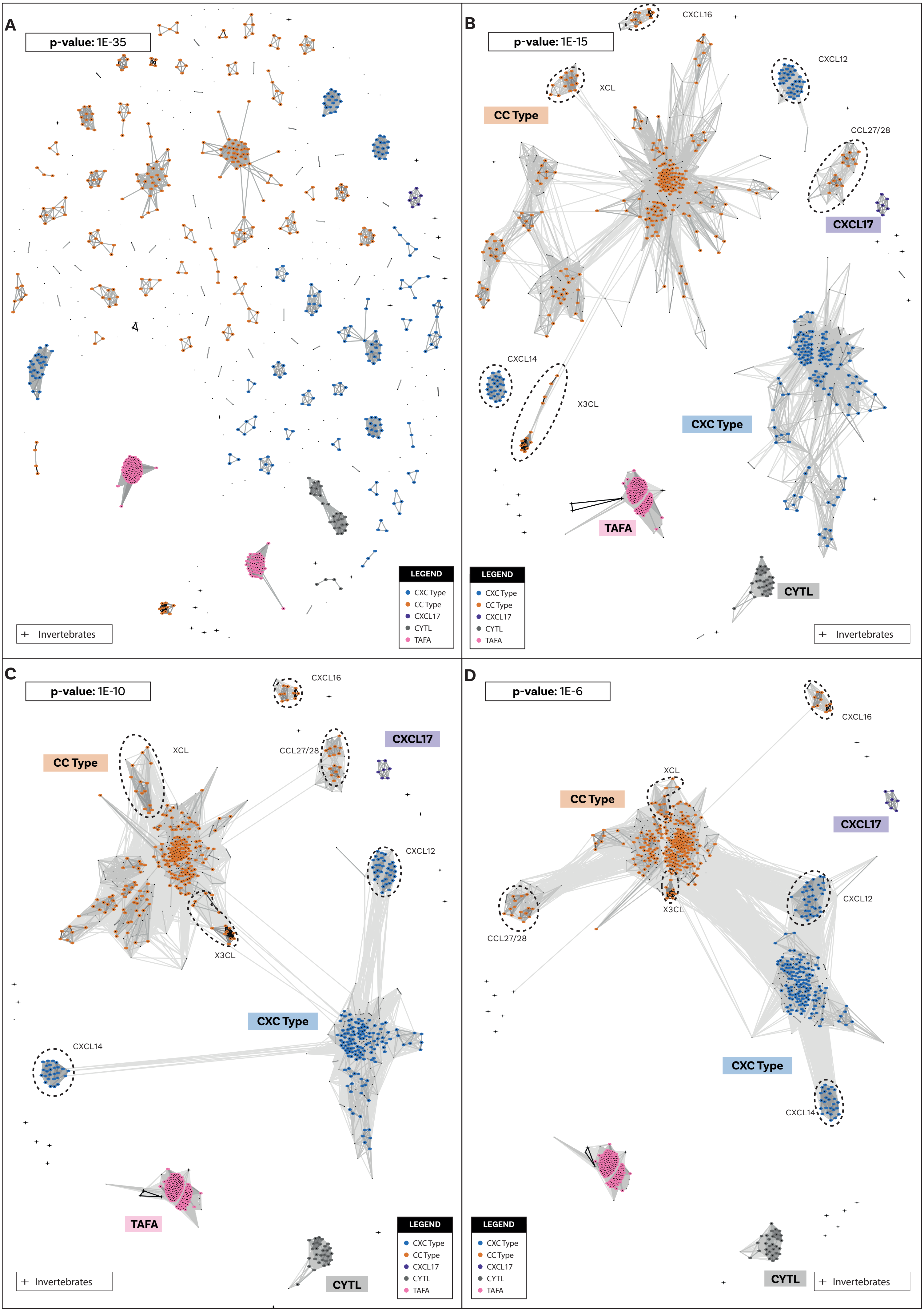
A
p-value: 1E-35
Invertebrates
LEGEND
CXC Type
CC Type
CXCL17
CYTL
TAFA
B
p-value: 1E-15
CXCL16
XCL
CXCL12
CC Type
CCL27/28
CXCL17
CXCL14
X3CL
CXC Type
TAFA
CYTL
LEGEND
CXC Type
CC Type
CXCL17
CYTL
TAFA
Invertebrates
C
p-value: 1E-10
CXCL16
XCL
CC Type
CCL27/28
CXCL17
CXCL12
X3CL
CXC Type
CXCL14
TAFA
CYTL
LEGEND
CXC Type
CC Type
CXCL17
CYTL
TAFA
Invertebrates
D
p-value: 1E-6
CXCL16
XCL
CC Type
CXCL17
CXCL12
X3CL
CCL27/28
CXC Type
CXCL14
CYTL
LEGEND
CXC Type
CC Type
CXCL17
CYTL
TAFA
Invertebrates

Fig. S2

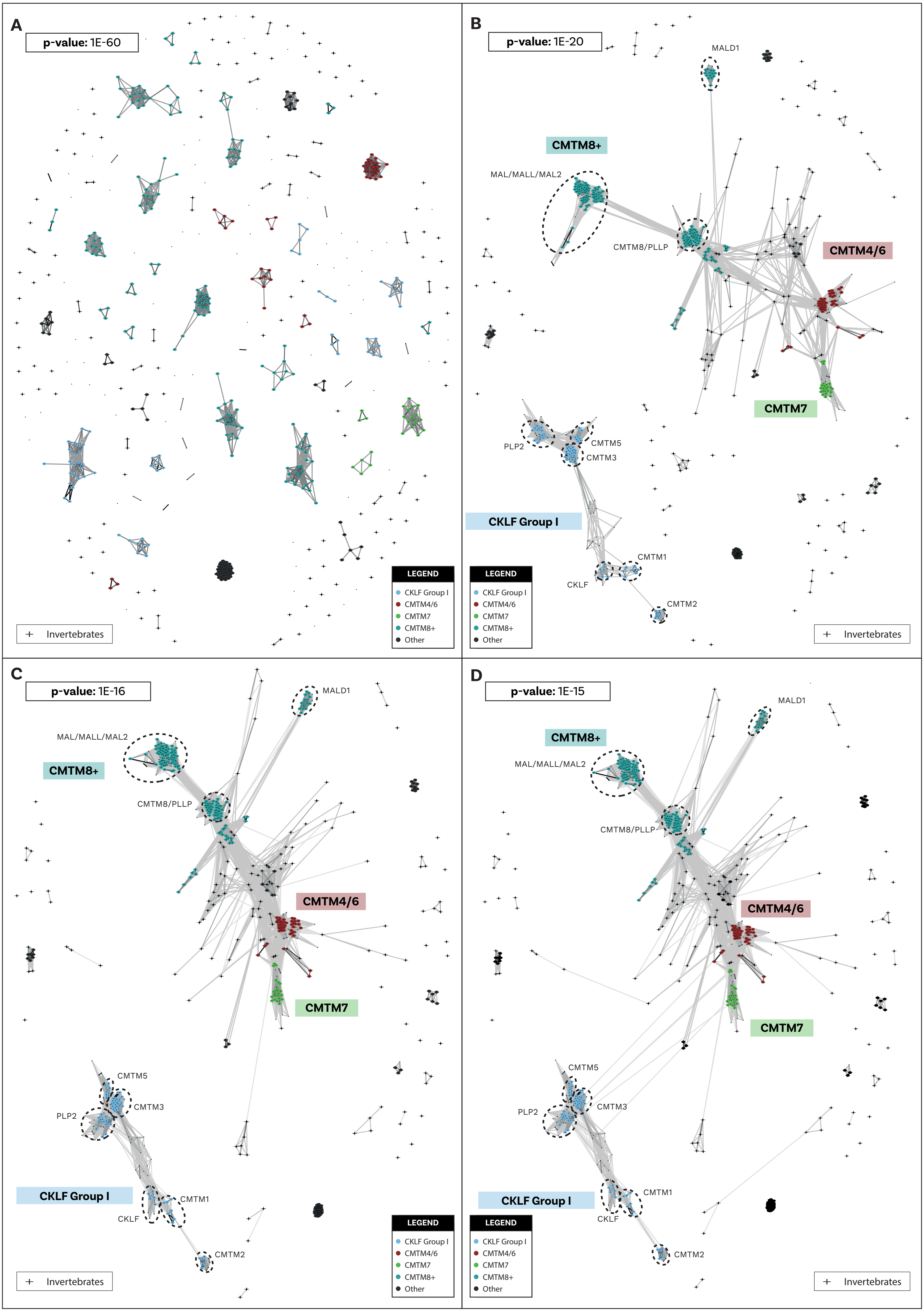
A
p-value: 1E-60
LEGEND
CKLF Group I
CMTM4/6
CMTM7
CMTM8+
Other
+ Invertebrates
B
p-value: 1E-20
MALD1
CMTM8+
MAL/MALL/MAL2
CMTM8/PLLP
CMTM4/6
CMTM7
PLP2
CMTM5
CMTM3
CKLF Group I
CMTM1
CKLF
CMTM2
LEGEND
CKLF Group I
CMTM4/6
CMTM7
CMTM8+
Other
+ Invertebrates
C
p-value: 1E-16
MALD1
MAL/MALL/MAL2
CMTM8+
CMTM8/PLLP
CMTM4/6
CMTM7
CMTM5
CMTM3
PLP2
CKLF Group I
CMTM1
CKLF
CMTM2
LEGEND
CKLF Group I
CMTM4/6
CMTM7
CMTM8+
Other
+ Invertebrates
D
p-value: 1E-15
MALD1
CMTM8+
MAL/MALL/MAL2
CMTM8/PLLP
CMTM4/6
CMTM7
CMTM5
PLP2
CMTM3
CKLF Group I
CMTM1
CKLF
CMTM2
LEGEND
CKLF Group I
CMTM4/6
CMTM7
CMTM8+
Other
+ Invertebrates

Fig. S3

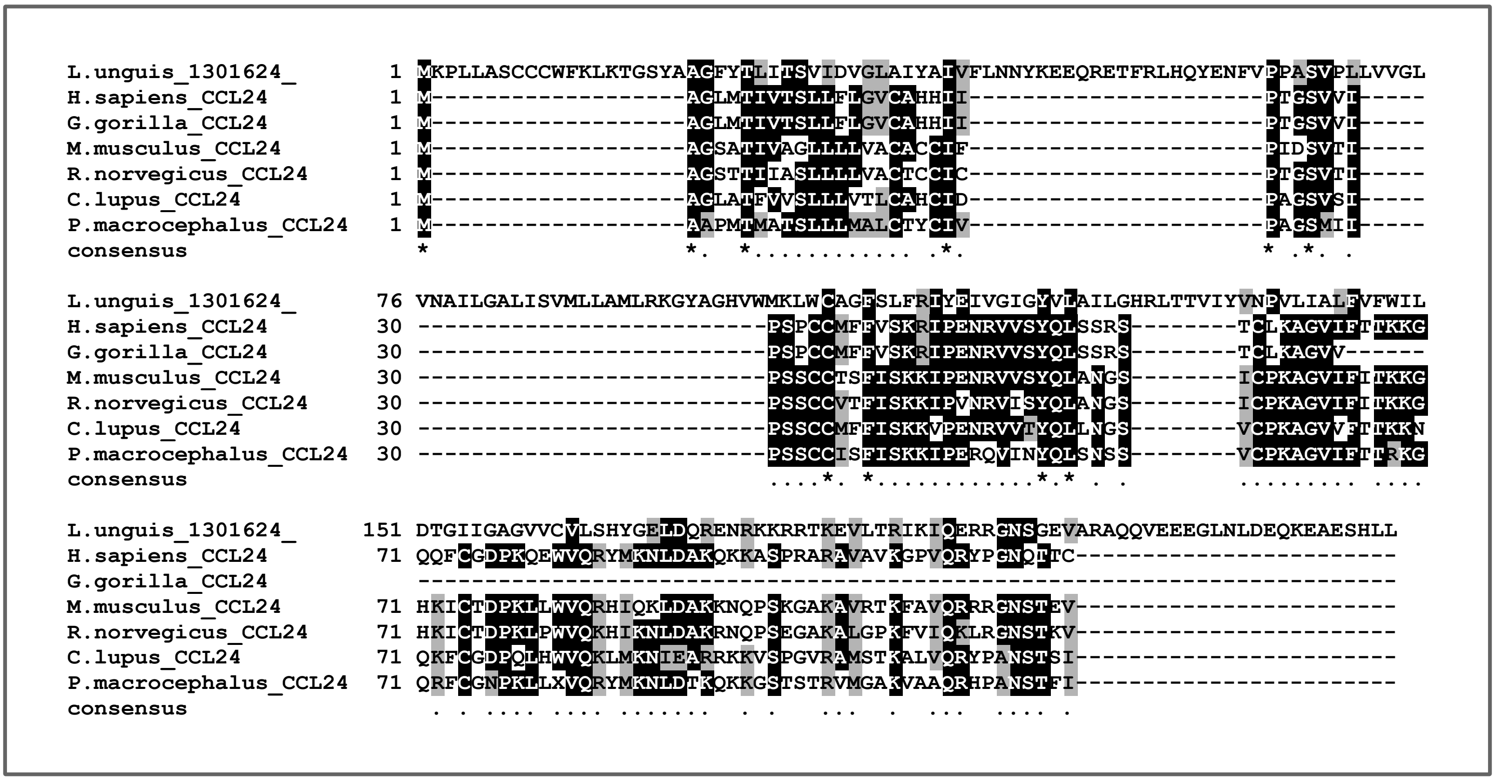
L.unguis_1301624_ 1 MKPLLASCCCWFKLKTGSYAAGFYTLITSVIDVGLAIYAIVFLNNYKEEQRETFRLHQYENFVPPASVPLLVVGL
H.sapiens_CCL24 1 M-------------------AGLMTIVTSLLFLGVCAHHII----------------------PTGSVVI-----
G.gorilla_CCL24 1 M-------------------AGLMTIVTSLLFLGVCAHHII----------------------PTGSVVI-----
M.musculus_CCL24 1 M-------------------AGSATIVAGLLLLVACACCIF----------------------PIDSVTI-----
R.norvegicus_CCL24 1 M-------------------AGSTTIIASLLLLVACTCCIC----------------------PTGSVTI-----
C.lupus_CCL24 1 M-------------------AGLATFVVSLLLVTLCAHCID----------------------PAGSVSI-----
P.macrocephalus_CCL24 1 M-------------------AAPMTMATSLLLMALCTYCIV----------------------PAGSMII-----
consensus *                   *.  *............ .*.                      * .*. .
L.unguis_1301624_ 76 VNAILGALISVMLLAMLRKGYAGHVWMKLWCAGFSLFRIYEIVGIGYVLAILGHRLTTVIYVNPVLIALFVFWIL
H.sapiens_CCL24 30 --------------------------PSPCCMFFVSKRIPENRVVSYQLSSRS--------TCLKAGVIFTTKKG
G.gorilla_CCL24 30 --------------------------PSPCCMFFVSKRIPENRVVSYQLSSRS--------TCLKAGVV------
M.musculus_CCL24 30 --------------------------PSSCCTSFISKKIPENRVVSYQLANGS--------ICPKAGVIFITKKG
R.norvegicus_CCL24 30 --------------------------PSSCCVTFISKKIPVNRVISYQLANGS--------ICPKAGVIFITKKG
C.lupus_CCL24 30 --------------------------PSSCCMFFISKKVPENRVVTYQLLNGS--------VCPKAGVVFTTKKN
P.macrocephalus_CCL24 30 --------------------------PSSCCISFISKKIPERQVINYQLSNSS--------VCPKAGVIFTTRKG
consensus ....*. *...........*.* . .        ......... ....
L.unguis_1301624_ 151 DTGIIGAGVVCVLSHYGELDQRENRKKRRTKEVLTRIKIQERRGNSGEVARAQQVEEEGLNLDEQKEAESHLL
H.sapiens_CCL24 71 QQFCGDPKQEWVQRYMKNLDAKQKKASPRARAVAVKGPVQRYPGNQTTC------------------------
G.gorilla_CCL24 -------------------------------------------------------------------------
M.musculus_CCL24 71 HKICTDPKLLWVQRHIQKLDAKKNQPSKGAKAVRTKFAVQRRRGNSTEV------------------------
R.norvegicus_CCL24 71 HKICTDPKLPWVQKHIKNLDAKRNQPSEGAKALGPKFVIQKLRGNSTKV------------------------
C.lupus_CCL24 71 QKFCGDPQLHWVQKLMKNIEARRKKVSPGVRAMSTKALVQRYPANSTSI------------------------
P.macrocephalus_CCL24 71 QRFCGNPKLLXVQRYMKNLDTKQKKGSTSTRVMGAKVAAQRHPANSTFI------------------------
consensus . . .... .... ....... . .   ...  .  ...  .... .

**Fig. S4**

```
C.hemisphaerica_3093647_    1 LQMDYSVKYIFKQNDIGDDGNVLLFHYGDAYYLLHWFGCCLQFFKLKKVDEQQQQSLQEINASRKNQQFTPDYKY
H.sapiens_CCL3              1 MQ-------------------------------------------------------------------------
G.gorilla_CCL3              1 MQ-------------------------------------------------------------------------
M.musculus_CCL3             1 MK-------------------------------------------------------------------------
R.norvegicus_CCL3           1 MK-------------------------------------------------------------------------
C.lupus_CCL3                1 ME-------------------------------------------------------------------------
P.macrocephalus_CL3L        1 MK-------------------------------------------------------------------------
consensus                     .

C.hemisphaerica_3093647_   76 SLKERIRLIVFNDVTNSIIILNLKGGISITAAEKVLKIQQNDEEMDYEPGDIFNNLLAQSVQPSNELNLESIVTV
H.sapiens_CCL3              3 ---------------------------------------------------------------------------
G.gorilla_CCL3              3 ---------------------------------------------------------------------------
M.musculus_CCL3             3 ---------------------------------------------------------------------------
R.norvegicus_CCL3           3 ---------------------------------------------------------------------------
C.lupus_CCL3                3 ---------------------------------------------------------------------------
P.macrocephalus_CL3L        3 ---------------------------------------------------------------------------
consensus

C.hemisphaerica_3093647_  151 SNVFTIKTVENYLYVISSASKSNILTRIYLIERNEDGLRLSPKFKQNLGGEIRSDAIC-------LPENFTA-YQ
H.sapiens_CCL3              3 ----------------VSTAALAVLLCTMALCNQ---------FSASLAADTPT-ACCFSYTSRQIPQNFIADYF
G.gorilla_CCL3              3 ----------------VSTAALAVLLCTMALCNQ---------FSASLAADMPT-ACCFSYISRQIPQNFIADYF
M.musculus_CCL3             3 ----------------VSTTALAVLLCTMTLCNQ--------VFSAPYGADTPT-ACCFSY-SRKIPRQFIVDYF
R.norvegicus_CCL3           3 ----------------VSTAALAVLLCTMALWNE--------VFSAPYGADTPT-ACCFSY-GRQIPRKFIADYF
C.lupus_CCL3                3 ----------------VSTAALAVLLLIAALCSQ--------TCSSTFGADTPT-SCCFSYISRQIPRKFVVDYY
P.macrocephalus_CL3L        3 ----------------VPAAALAVLLCPVALCSQ--------VFSAPFETDTLT-ACCFSYTAWQLPRKFVADYF
consensus                                     .........*....*...         ... ....... ..*... ....*..*...*.

C.hemisphaerica_3093647_  218 ELNSNLELPGIIFSTSNGLILCSNPSSA-TPSLIYDLKQRLAGVVHVTLPKTTELSTNNLIPSSQPKDVSLNGDC
H.sapiens_CCL3             52 ETSSQCSKPGVIFLTKRSRQVCADPSEEWVQKYVSDL--------------------------------------
G.gorilla_CCL3             52 ETSSQCSKPGVIFLTKRSRQVCADPSEEWVQKYVSDL--------------------------------------
M.musculus_CCL3            52 ETSSLCSQPGVIFLTKRNRQICADSKETWVQEYITDL--------------------------------------
R.norvegicus_CCL3          52 ETSSLCSQPGVIFLTKRNRQICADPKETWVQEYITEL--------------------------------------
C.lupus_CCL3               53 ETSSQCSKPGVIFQTKRGQQICADPGKAWVQKYITNL--------------------------------------
P.macrocephalus_CL3L       53 ETSSQCSKPGVIFQTKRGRQLCANPSEDWVQEYITDL--------------------------------------
consensus                     *..*....**.**.*.. ...*..... ... ....*

C.hemisphaerica_3093647_  292 LCFVGENGRVSVAIPNNTIKRKCIEFTVGQLSINVVFCTSFQDILIAASEDHILFFKLVCKCKIESKHSPNRKII
H.sapiens_CCL3             89 ---------------------------------------------------------------------------
G.gorilla_CCL3             89 ---------------------------------------------------------------------------
M.musculus_CCL3            89 ---------------------------------------------------------------------------
R.norvegicus_CCL3          89 ---------------------------------------------------------------------------
C.lupus_CCL3               90 ---------------------------------------------------------------------------
P.macrocephalus_CL3L       90 ---------------------------------------------------------------------------
consensus

C.hemisphaerica_3093647_  367 NVSFSLLKEIKIPNVNKLIMDPLNPGVIICHNTAKSIMVVNLGDFWVTKEGDCQKDPRTVTSLLKVYEEQNKENE
H.sapiens_CCL3             89 ---------------------------------------------------------------------------
G.gorilla_CCL3             89 ---------------------------------------------------------------------------
M.musculus_CCL3            89 ---------------------------------------------------------------------------
R.norvegicus_CCL3          89 ---------------------------------------------------------------------------
C.lupus_CCL3               90 ---------------------------------------------------------------------------
P.macrocephalus_CL3L       90 ---------------------------------------------------------------------------
consensus

C.hemisphaerica_3093647_  442 LLKNESVELNKTLIHLSIVGEWVKSIKKDSKSCFGIDKRITVVRSNSKKLAFLTLSIQNKSKLAVPAFCSINIQY
H.sapiens_CCL3             89 -------ELSA----------------------------------------------------------------
G.gorilla_CCL3             89 -------ELSA----------------------------------------------------------------
M.musculus_CCL3            89 -------ELNA----------------------------------------------------------------
R.norvegicus_CCL3          89 -------ELNA----------------------------------------------------------------
C.lupus_CCL3               90 -------ELNA----------------------------------------------------------------
P.macrocephalus_CL3L       90 -------QLNA----------------------------------------------------------------
consensus                            .*..

C.hemisphaerica_3093647_  517 QSQFQPKMIKTETIQLNTSPPTAKQSITIELSDRFMRFSPIDFNTSITFSLQTYQSISLPLRTFTVNILDFCDIS
H.sapiens_CCL3                ---------------------------------------------------------------------------
G.gorilla_CCL3                ---------------------------------------------------------------------------
M.musculus_CCL3               ---------------------------------------------------------------------------
R.norvegicus_CCL3             ---------------------------------------------------------------------------
C.lupus_CCL3                  ---------------------------------------------------------------------------
P.macrocephalus_CL3L          ---------------------------------------------------------------------------
consensus

C.hemisphaerica_3093647_  592 PTMFSMSKQMLAFPKTYLEQQNQPSHSICIKLDPTKFDNNKDAFCNCLSTCLNIEIKKTKENFLRYNIEYIHGET
H.sapiens_CCL3                ---------------------------------------------------------------------------
G.gorilla_CCL3                ---------------------------------------------------------------------------
M.musculus_CCL3               ---------------------------------------------------------------------------
R.norvegicus_CCL3             ---------------------------------------------------------------------------
C.lupus_CCL3                  ---------------------------------------------------------------------------
P.macrocephalus_CL3L          ---------------------------------------------------------------------------
consensus

C.hemisphaerica_3093647_  667 AEVEIKLDNFIVSVSTKNVMFLASLHHHFSGCLKEFTILNGECGEETKQKIVESLKELSSAFTKNITGQDSADVI
H.sapiens_CCL3                ---------------------------------------------------------------------------
G.gorilla_CCL3                ---------------------------------------------------------------------------
M.musculus_CCL3               ---------------------------------------------------------------------------
R.norvegicus_CCL3             ---------------------------------------------------------------------------
C.lupus_CCL3                  ---------------------------------------------------------------------------
P.macrocephalus_CL3L          ---------------------------------------------------------------------------

C.hemisphaerica_3093647_  742 STLSKFYSLQRQ
H.sapiens_CCL3                ------------
G.gorilla_CCL3                ------------
M.musculus_CCL3               ------------
R.norvegicus_CCL3             ------------
C.lupus_CCL3                  ------------
P.macrocephalus_CL3L          ------------
consensus
```

**Fig. S5**

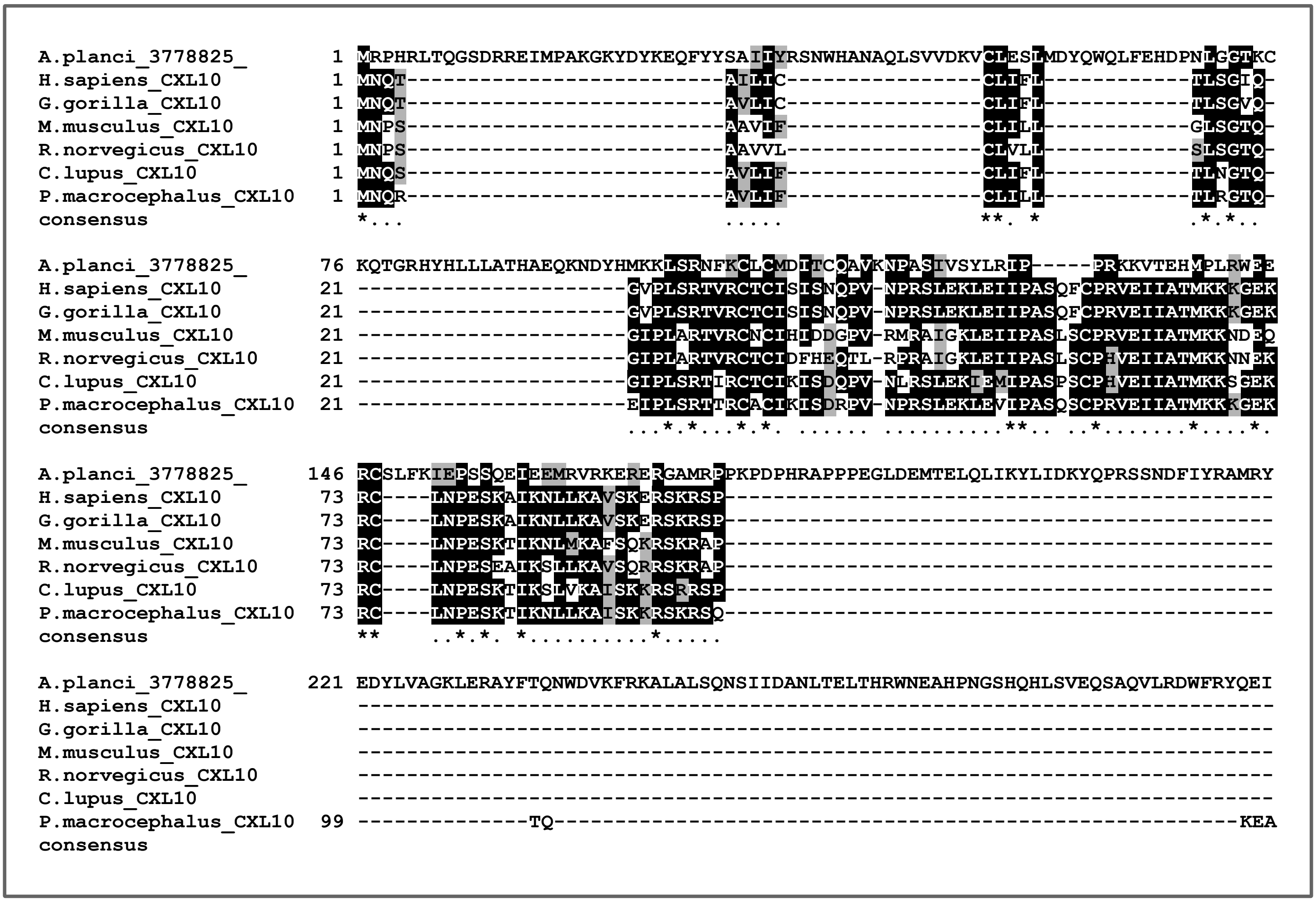

A.planci_3778825_ 1 MRPHRLTQGSDRREIMPAKGKYDYKEQFYYSAIIYRSNWHANAQLSVVDKVCLESLMDYQWQLFEHDPNLGGTKC
H.sapiens_CXL10 1 MNQT--------------------------AILIC----------------CLIFL------------TLSGIQ-
G.gorilla_CXL10 1 MNQT--------------------------AVLIC----------------CLIFL------------TLSGVQ-
M.musculus_CXL10 1 MNPS--------------------------AAVIF----------------CLILL------------GLSGTQ-
R.norvegicus_CXL10 1 MNPS--------------------------AAVVL----------------CLVLL------------SLSGTQ-
C.lupus_CXL10 1 MNQS--------------------------AVLIF----------------CLIFL------------TLNGTQ-
P.macrocephalus_CXL10 1 MNQR--------------------------AVLIF----------------CLILL------------TLRGTQ-
consensus *... ..... **. * .*.*..
A.planci_3778825_ 76 KQTGRHYHLLLATHAEQKNDYHMKKLSRNFKCLCMDITCQAVKNPASIVSYLRIP-----PRKKVTEHMPLRWEE
H.sapiens_CXL10 21 ----------------------GVPLSRTVRCTCISISNQPV-NPRSLEKLEIIPASQFCPRVEIIATMKKKGEK
G.gorilla_CXL10 21 ----------------------GVPLSRTVRCTCISISNQPV-NPRSLEKLEIIPASQFCPRVEIIATMKKKGEK
M.musculus_CXL10 21 ----------------------GIPLARTVRCNCIHIDDGPV-RMRAIGKLEIIPASLSCPRVEIIATMKKNDEQ
R.norvegicus_CXL10 21 ----------------------GIPLARTVRCTCIDFHEQTL-RPRAIGKLEIIPASLSCPHVEIIATMKKNNEK
C.lupus_CXL10 21 ----------------------GIPLSRTIRCTCIKISDQPV-NLRSLEKIEMIPASPSCPHVEIIATMKKSGEK
P.macrocephalus_CXL10 21 ----------------------EIPLSRTTRCACIKISDRPV-NPRSLEKLEVIPASQSCPRVEIIATMKKKGEK
consensus ...*.*...*.*. ...... ..........**.. ..*.......*....*.
A.planci_3778825_ 146 RCSLFKIEPSSQEIEEMRVRKERERGAMRPPKPDPHRAPPPEGLDEMTELQLIKYLIDKYQPRSSNDFIYRAMRY
H.sapiens_CXL10 73 RC----LNPESKAIKNLLKAVSKERSKRSP---------------------------------------------
G.gorilla_CXL10 73 RC----LNPESKAIKNLLKAVSKERSKRSP---------------------------------------------
M.musculus_CXL10 73 RC----LNPESKTIKNLMKAFSQKRSKRAP---------------------------------------------
R.norvegicus_CXL10 73 RC----LNPESEAIKSLLKAVSQRRSKRAP---------------------------------------------
C.lupus_CXL10 73 RC----LNPESKTIKSLVKAISKKRSRRSP---------------------------------------------
P.macrocephalus_CXL10 73 RC----LNPESKTIKNLLKAISKKRSKRSQ---------------------------------------------
consensus ** ..*.*. *.........*.....
A.planci_3778825_ 221 EDYLVAGKLERAYFTQNWDVKFRKALALSQNSIIDANLTELTHRWNEAHPNGSHQHLSVEQSAQVLRDWFRYQEI
H.sapiens_CXL10 ---------------------------------------------------------------------------
G.gorilla_CXL10 ---------------------------------------------------------------------------
M.musculus_CXL10 ---------------------------------------------------------------------------
R.norvegicus_CXL10 ---------------------------------------------------------------------------
C.lupus_CXL10 ---------------------------------------------------------------------------
P.macrocephalus_CXL10 99 --------------TQ--------------------------------------------------------KEA
consensus

**Fig. S6**

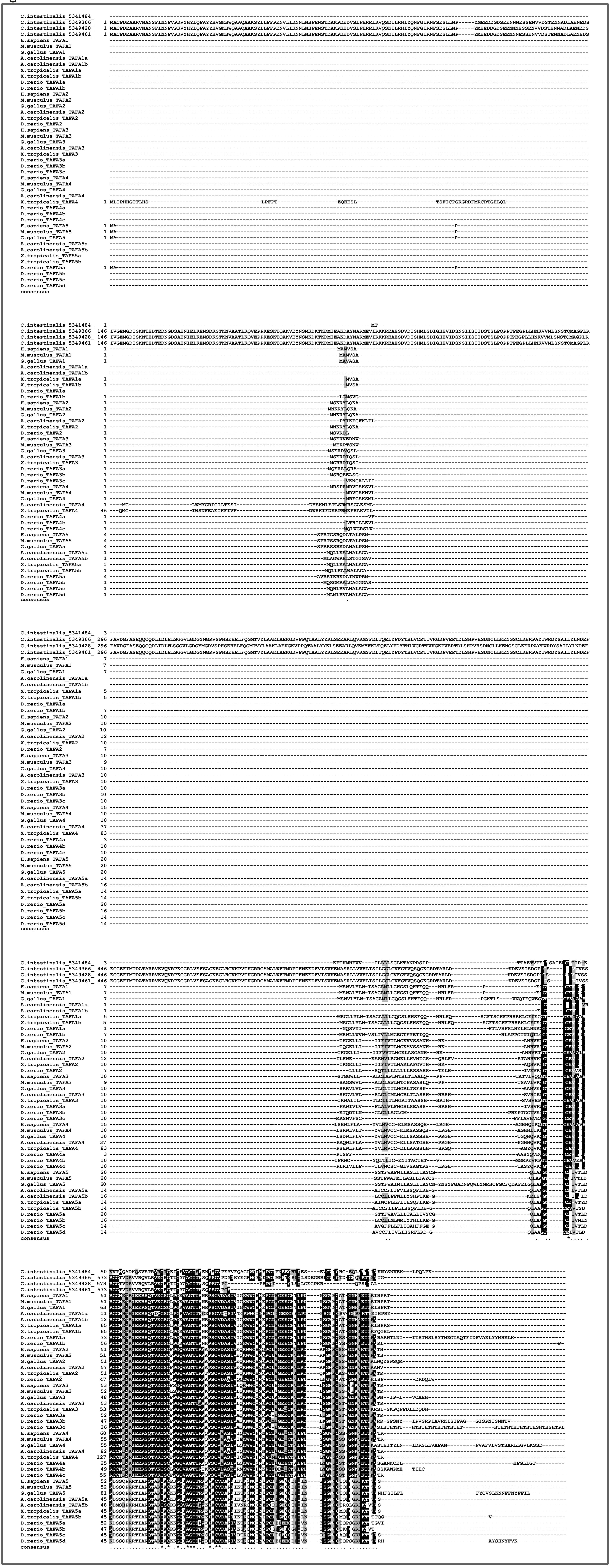

**Fig. S7**

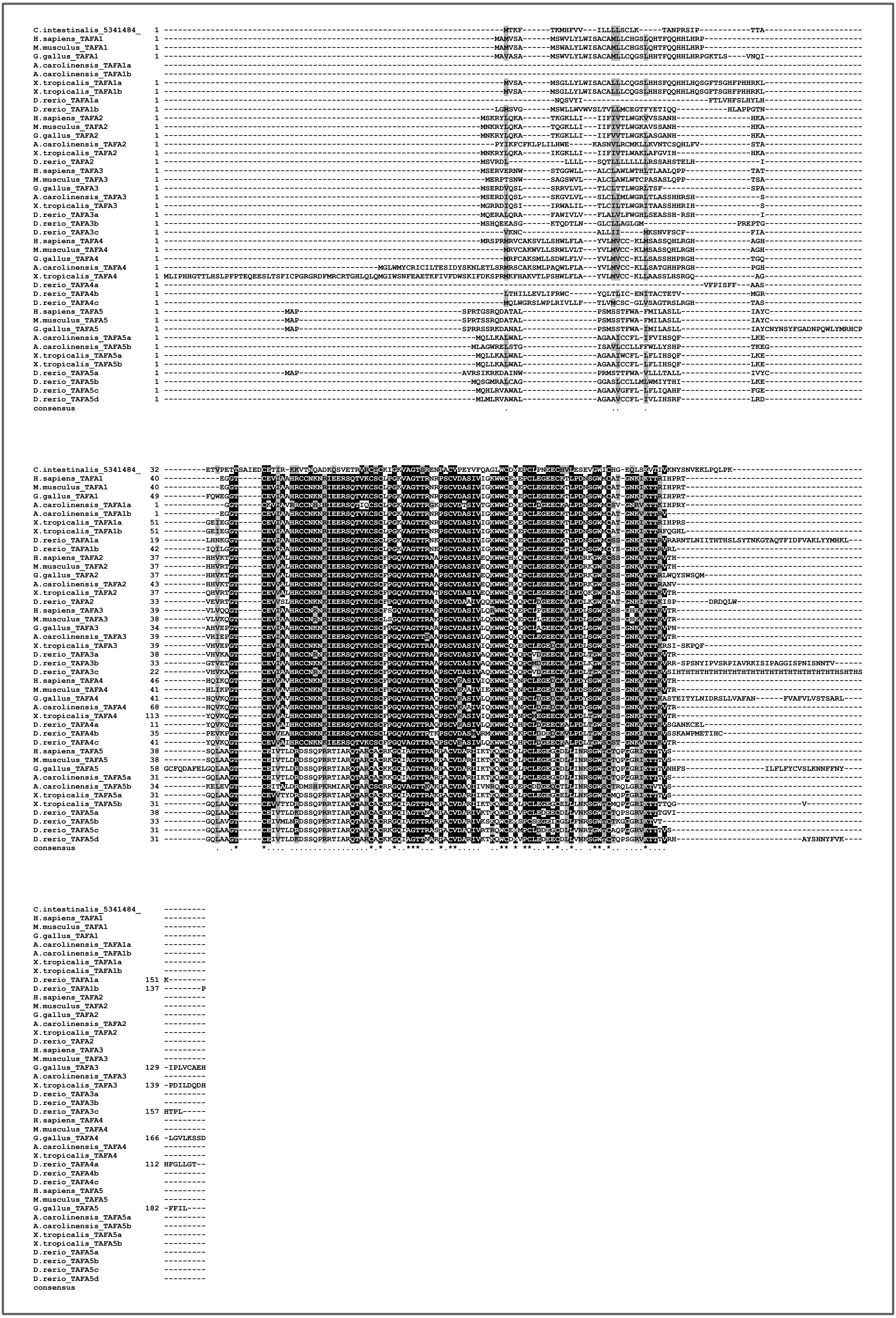
C.intestinalis_5341484_
H.sapiens_TAFA1
M.musculus_TAFA1
G.gallus_TAFA1
A.carolinensis_TAFA1a
A.carolinensis_TAFA1b
X.tropicalis_TAFA1a
X.tropicalis_TAFA1b
D.rerio_TAFA1a
D.rerio_TAFA1b
H.sapiens_TAFA2
M.musculus_TAFA2
G.gallus_TAFA2
A.carolinensis_TAFA2
X.tropicalis_TAFA2
D.rerio_TAFA2
H.sapiens_TAFA3
M.musculus_TAFA3
G.gallus_TAFA3
A.carolinensis_TAFA3
X.tropicalis_TAFA3
D.rerio_TAFA3a
D.rerio_TAFA3b
D.rerio_TAFA3c
H.sapiens_TAFA4
M.musculus_TAFA4
G.gallus_TAFA4
A.carolinensis_TAFA4
X.tropicalis_TAFA4
D.rerio_TAFA4a
D.rerio_TAFA4b
D.rerio_TAFA4c
H.sapiens_TAFA5
M.musculus_TAFA5
G.gallus_TAFA5
A.carolinensis_TAFA5a
A.carolinensis_TAFA5b
X.tropicalis_TAFA5a
X.tropicalis_TAFA5b
D.rerio_TAFA5a
D.rerio_TAFA5b
D.rerio_TAFA5c
D.rerio_TAFA5d
consensus

Fig. S8

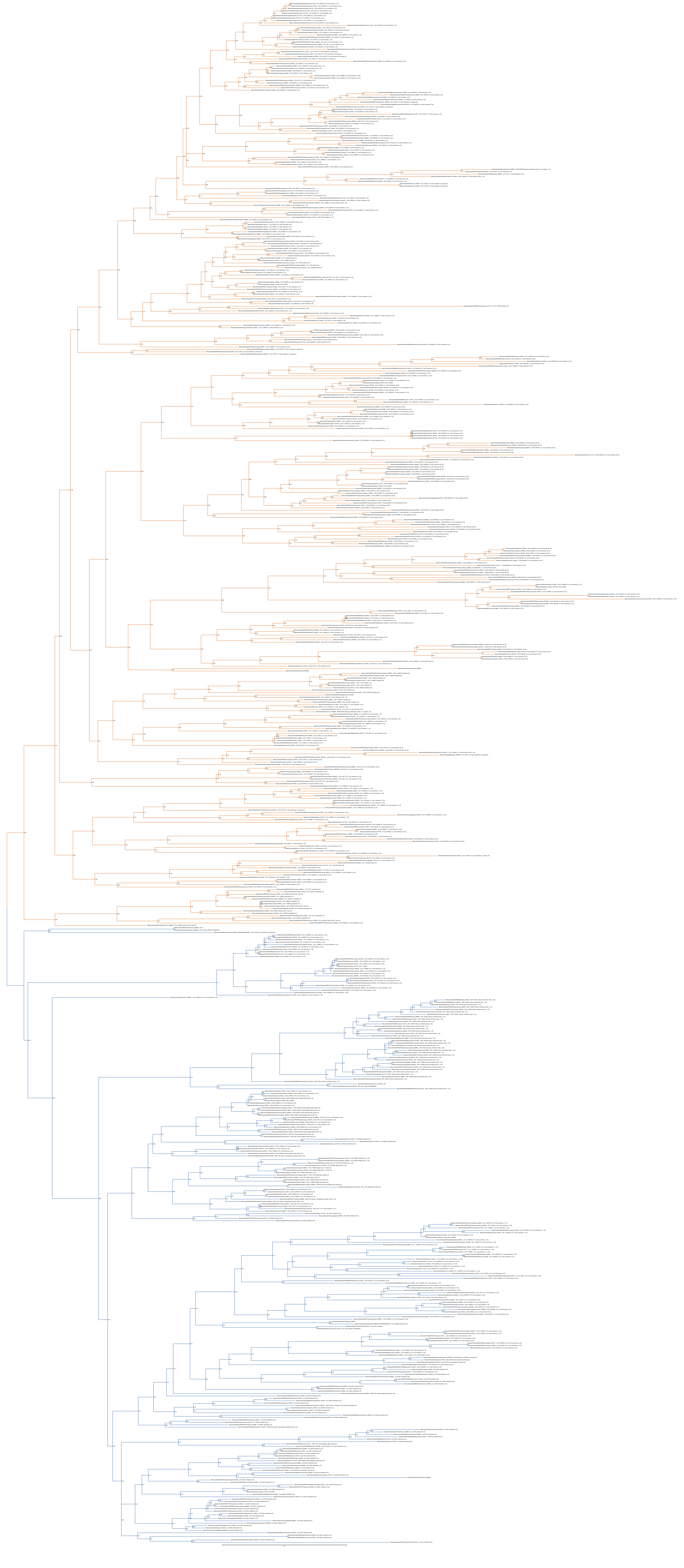

Fig. S9

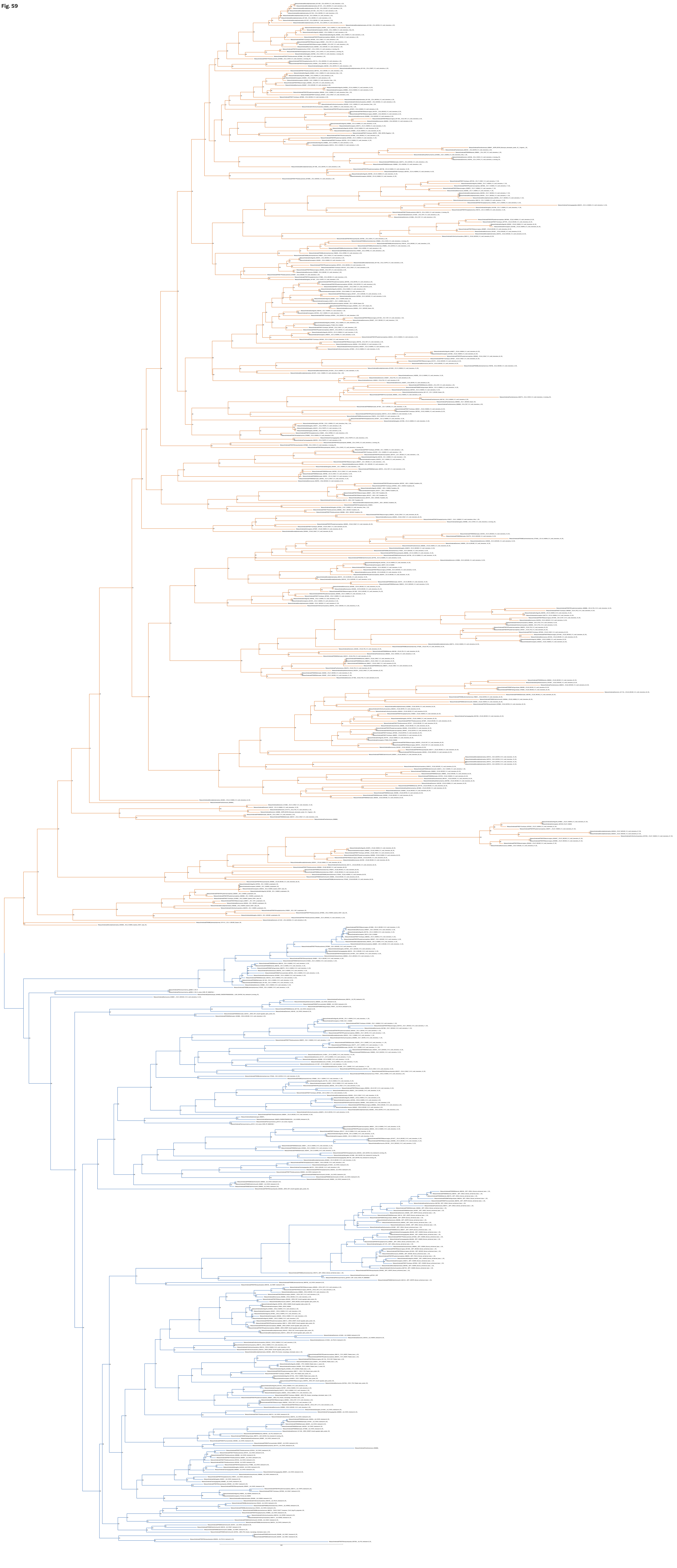

**Fig. S10**

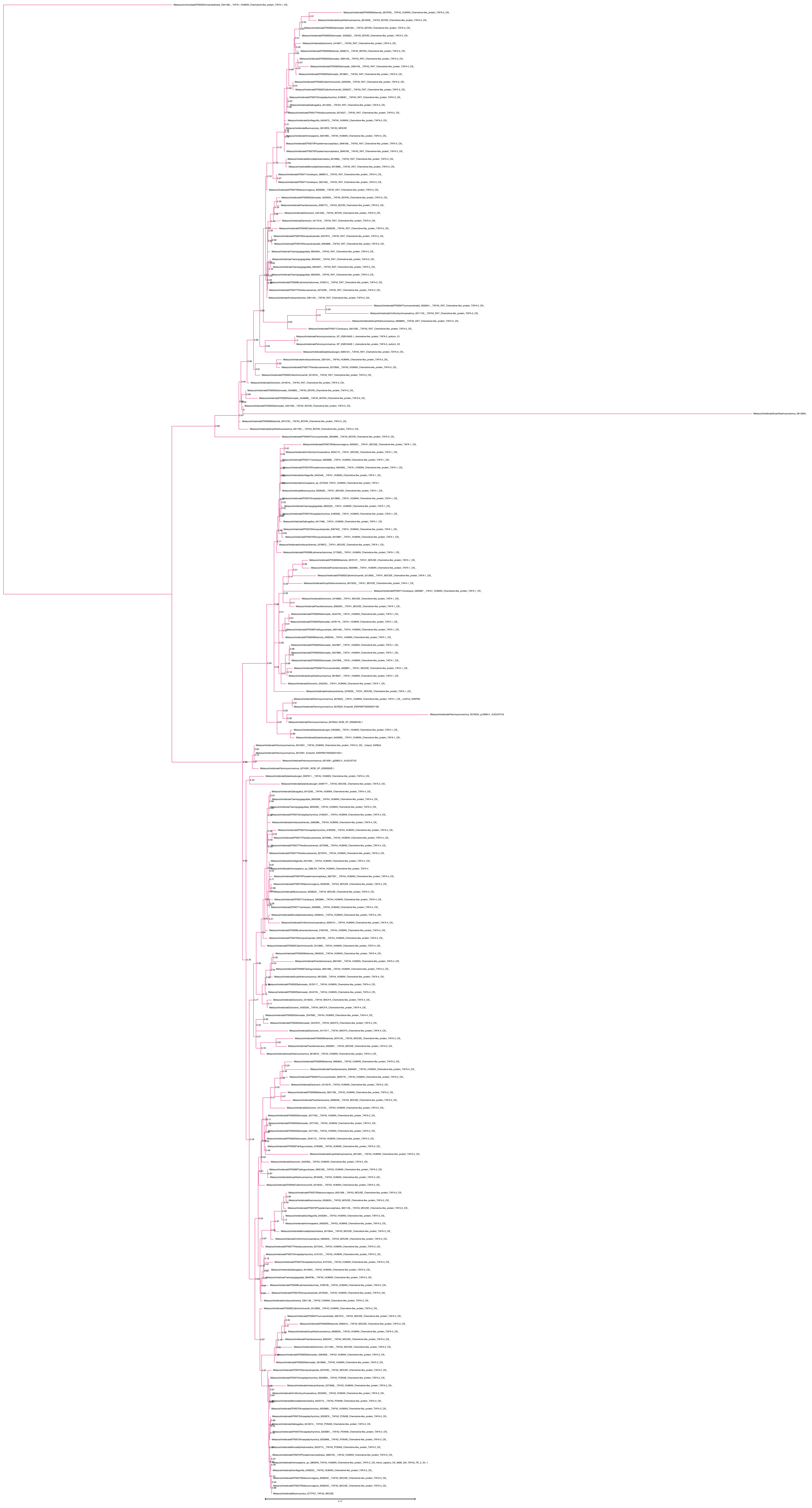

**Fig. S11**

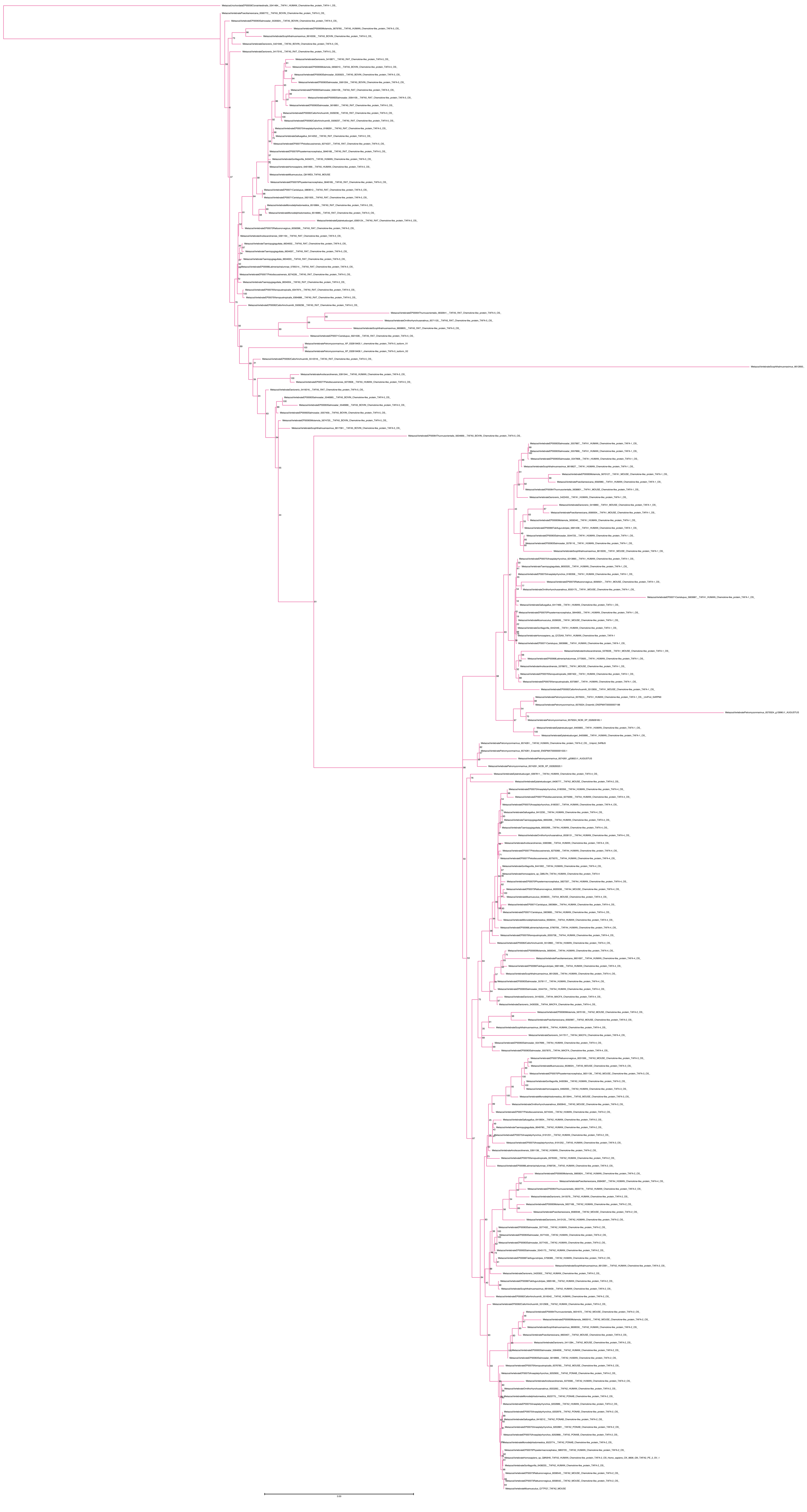

**Fig. S12**

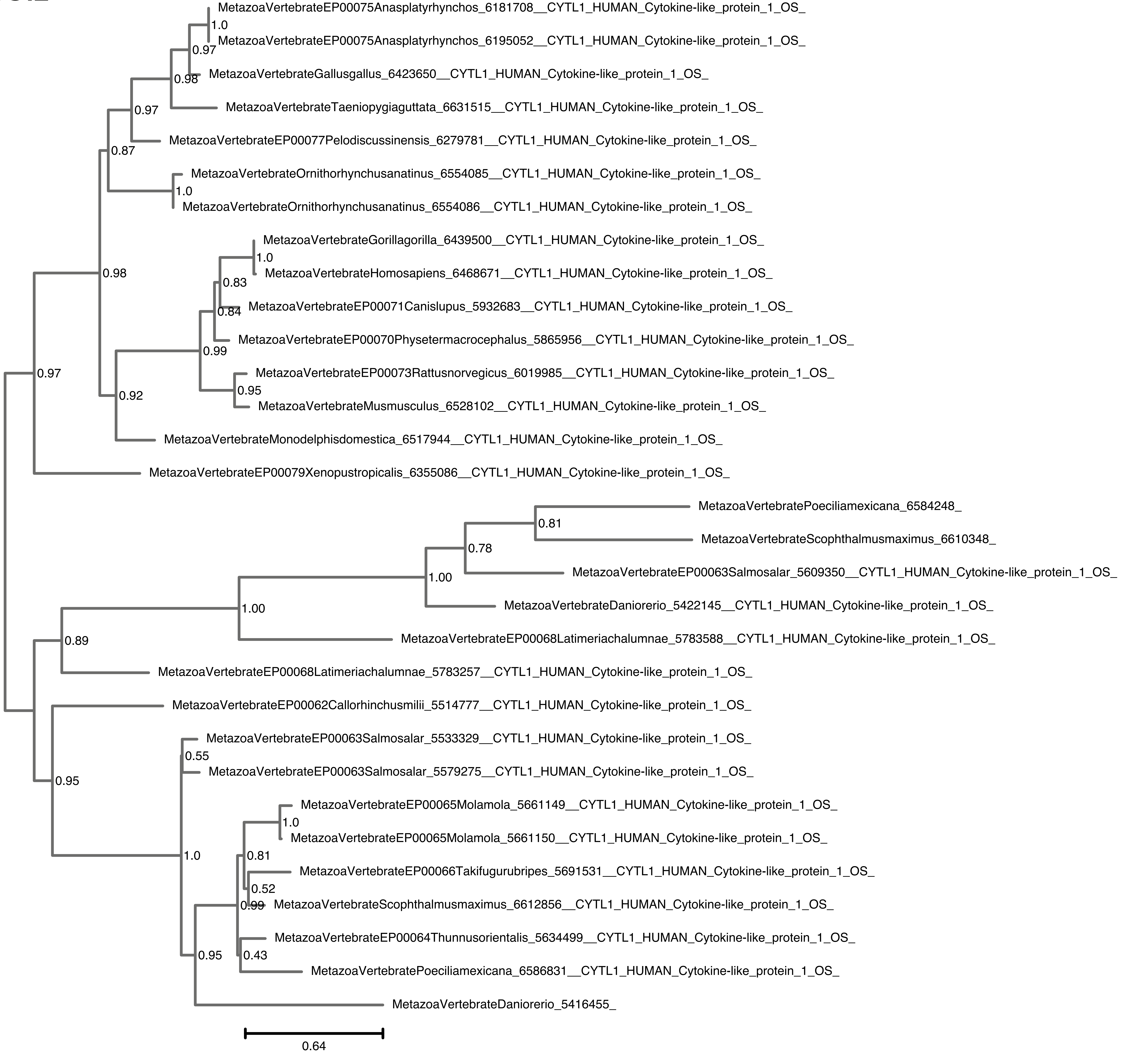
MetazoaVertebrateEP00075Anasplatyrhynchos_6181708__CYTL1_HUMAN_Cytokine-like_protein_1_OS_
1.0
MetazoaVertebrateEP00075Anasplatyrhynchos_6195052__CYTL1_HUMAN_Cytokine-like_protein_1_OS_
0.97
MetazoaVertebrateGallusgallus_6423650__CYTL1_HUMAN_Cytokine-like_protein_1_OS_
0.98
MetazoaVertebrateTaeniopygiaguttata_6631515__CYTL1_HUMAN_Cytokine-like_protein_1_OS_
0.97
MetazoaVertebrateEP00077Pelodiscussinensis_6279781__CYTL1_HUMAN_Cytokine-like_protein_1_OS_
0.87
MetazoaVertebrateOrnithorhynchusanatinus_6554085__CYTL1_HUMAN_Cytokine-like_protein_1_OS_
1.0
MetazoaVertebrateOrnithorhynchusanatinus_6554086__CYTL1_HUMAN_Cytokine-like_protein_1_OS_
MetazoaVertebrateGorillagorilla_6439500__CYTL1_HUMAN_Cytokine-like_protein_1_OS_
1.0
0.98
MetazoaVertebrateHomosapiens_6468671__CYTL1_HUMAN_Cytokine-like_protein_1_OS_
0.83
MetazoaVertebrateEP00071Canislupus_5932683__CYTL1_HUMAN_Cytokine-like_protein_1_OS_
0.84
MetazoaVertebrateEP00070Physetermacrocephalus_5865956__CYTL1_HUMAN_Cytokine-like_protein_1_OS_
0.99
MetazoaVertebrateEP00073Rattusnorvegicus_6019985__CYTL1_HUMAN_Cytokine-like_protein_1_OS_
0.97
0.95
0.92
MetazoaVertebrateMusmusculus_6528102__CYTL1_HUMAN_Cytokine-like_protein_1_OS_
MetazoaVertebrateMonodelphisdomestica_6517944__CYTL1_HUMAN_Cytokine-like_protein_1_OS_
MetazoaVertebrateEP00079Xenopustropicalis_6355086__CYTL1_HUMAN_Cytokine-like_protein_1_OS_
MetazoaVertebratePoeciliamexicana_6584248_
0.81
MetazoaVertebrateScophthalmusmaximus_6610348_
0.78
MetazoaVertebrateEP00063Salmosalar_5609350__CYTL1_HUMAN_Cytokine-like_protein_1_OS_
1.00
MetazoaVertebrateDaniorerio_5422145__CYTL1_HUMAN_Cytokine-like_protein_1_OS_
1.00
MetazoaVertebrateEP00068Latimeriachalumnae_5783588__CYTL1_HUMAN_Cytokine-like_protein_1_OS_
0.89
MetazoaVertebrateEP00068Latimeriachalumnae_5783257__CYTL1_HUMAN_Cytokine-like_protein_1_OS_
MetazoaVertebrateEP00062Callorhinchusmilii_5514777__CYTL1_HUMAN_Cytokine-like_protein_1_OS_
MetazoaVertebrateEP00063Salmosalar_5533329__CYTL1_HUMAN_Cytokine-like_protein_1_OS_
0.55
MetazoaVertebrateEP00063Salmosalar_5579275__CYTL1_HUMAN_Cytokine-like_protein_1_OS_
0.95
MetazoaVertebrateEP00065Molamola_5661149__CYTL1_HUMAN_Cytokine-like_protein_1_OS_
1.0
MetazoaVertebrateEP00065Molamola_5661150__CYTL1_HUMAN_Cytokine-like_protein_1_OS_
1.0
0.81
MetazoaVertebrateEP00066Takifugurubripes_5691531__CYTL1_HUMAN_Cytokine-like_protein_1_OS_
0.52
0.99
MetazoaVertebrateScophthalmusmaximus_6612856__CYTL1_HUMAN_Cytokine-like_protein_1_OS_
MetazoaVertebrateEP00064Thunnusorientalis_5634499__CYTL1_HUMAN_Cytokine-like_protein_1_OS_
0.43
0.95
MetazoaVertebratePoeciliamexicana_6586831__CYTL1_HUMAN_Cytokine-like_protein_1_OS_
MetazoaVertebrateDaniorerio_5416455_
0.64

**Fig. S13**

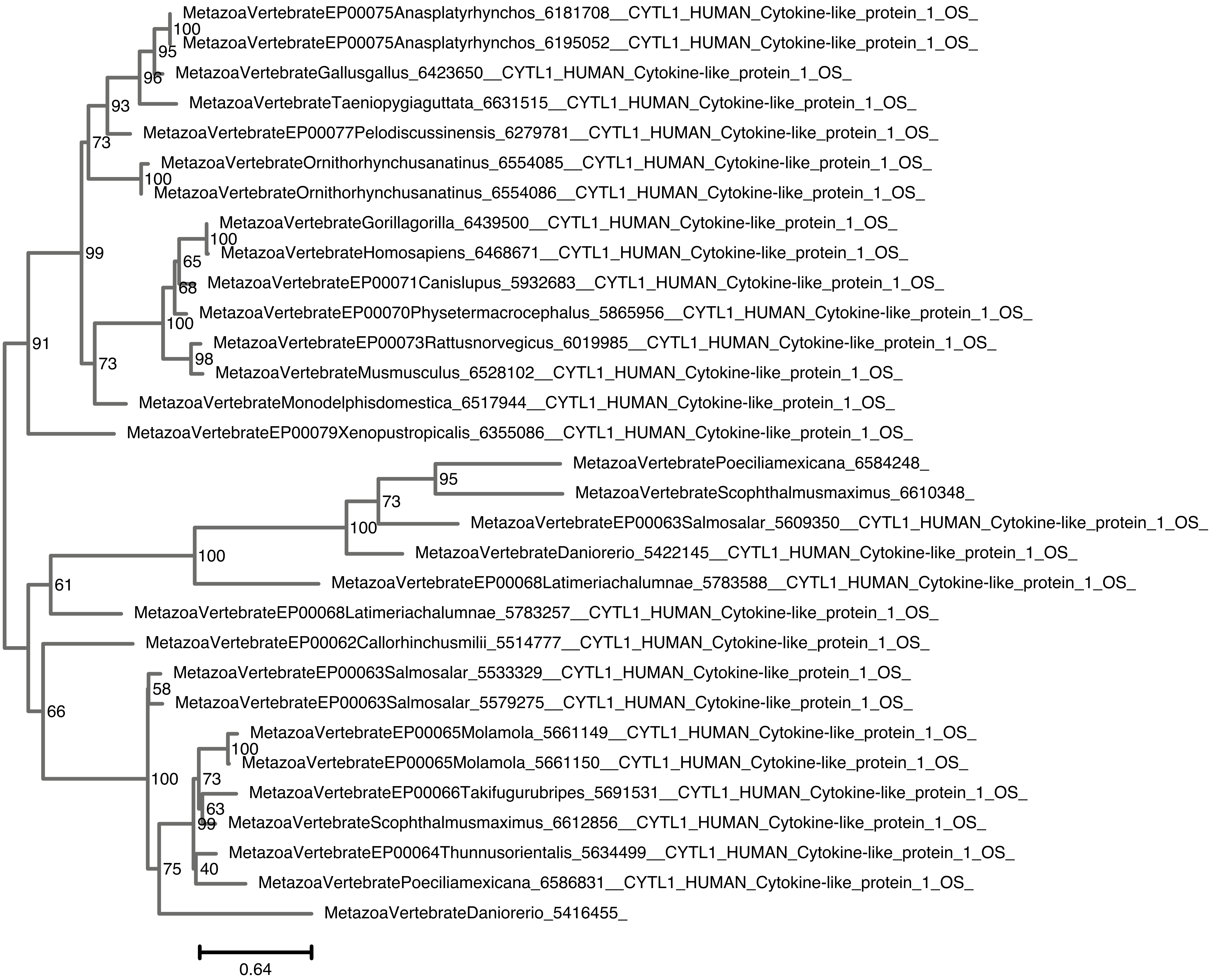
MetazoaVertebrateEP00075Anasplatyrhynchos_6181708__CYTL1_HUMAN_Cytokine-like_protein_1_OS_
MetazoaVertebrateEP00075Anasplatyrhynchos_6195052__CYTL1_HUMAN_Cytokine-like_protein_1_OS_
MetazoaVertebrateGallusgallus_6423650__CYTL1_HUMAN_Cytokine-like_protein_1_OS_
MetazoaVertebrateTaeniopygiaguttata_6631515__CYTL1_HUMAN_Cytokine-like_protein_1_OS_
MetazoaVertebrateEP00077Pelodiscussinensis_6279781__CYTL1_HUMAN_Cytokine-like_protein_1_OS_
MetazoaVertebrateOrnithorhynchusanatinus_6554085__CYTL1_HUMAN_Cytokine-like_protein_1_OS_
MetazoaVertebrateOrnithorhynchusanatinus_6554086__CYTL1_HUMAN_Cytokine-like_protein_1_OS_
MetazoaVertebrateGorillagorilla_6439500__CYTL1_HUMAN_Cytokine-like_protein_1_OS_
MetazoaVertebrateHomosapiens_6468671__CYTL1_HUMAN_Cytokine-like_protein_1_OS_
MetazoaVertebrateEP00071Canislupus_5932683__CYTL1_HUMAN_Cytokine-like_protein_1_OS_
MetazoaVertebrateEP00070Physetermacrocephalus_5865956__CYTL1_HUMAN_Cytokine-like_protein_1_OS_
MetazoaVertebrateEP00073Rattusnorvegicus_6019985__CYTL1_HUMAN_Cytokine-like_protein_1_OS_
MetazoaVertebrateMusmusculus_6528102__CYTL1_HUMAN_Cytokine-like_protein_1_OS_
MetazoaVertebrateMonodelphisdomestica_6517944__CYTL1_HUMAN_Cytokine-like_protein_1_OS_
MetazoaVertebrateEP00079Xenopustropicalis_6355086__CYTL1_HUMAN_Cytokine-like_protein_1_OS_
MetazoaVertebratePoeciliamexicana_6584248_
MetazoaVertebrateScophthalmusmaximus_6610348_
MetazoaVertebrateEP00063Salmosalar_5609350__CYTL1_HUMAN_Cytokine-like_protein_1_OS_
MetazoaVertebrateDaniorerio_5422145__CYTL1_HUMAN_Cytokine-like_protein_1_OS_
MetazoaVertebrateEP00068Latimeriachalumnae_5783588__CYTL1_HUMAN_Cytokine-like_protein_1_OS_
MetazoaVertebrateEP00068Latimeriachalumnae_5783257__CYTL1_HUMAN_Cytokine-like_protein_1_OS_
MetazoaVertebrateEP00062Callorhinchusmilii_5514777__CYTL1_HUMAN_Cytokine-like_protein_1_OS_
MetazoaVertebrateEP00063Salmosalar_5533329__CYTL1_HUMAN_Cytokine-like_protein_1_OS_
MetazoaVertebrateEP00063Salmosalar_5579275__CYTL1_HUMAN_Cytokine-like_protein_1_OS_
MetazoaVertebrateEP00065Molamola_5661149__CYTL1_HUMAN_Cytokine-like_protein_1_OS_
MetazoaVertebrateEP00065Molamola_5661150__CYTL1_HUMAN_Cytokine-like_protein_1_OS_
MetazoaVertebrateEP00066Takifugurubripes_5691531__CYTL1_HUMAN_Cytokine-like_protein_1_OS_
MetazoaVertebrateScophthalmusmaximus_6612856__CYTL1_HUMAN_Cytokine-like_protein_1_OS_
MetazoaVertebrateEP00064Thunnusorientalis_5634499__CYTL1_HUMAN_Cytokine-like_protein_1_OS_
MetazoaVertebratePoeciliamexicana_6586831__CYTL1_HUMAN_Cytokine-like_protein_1_OS_
MetazoaVertebrateDaniorerio_5416455_
0.64

**Fig. S14**

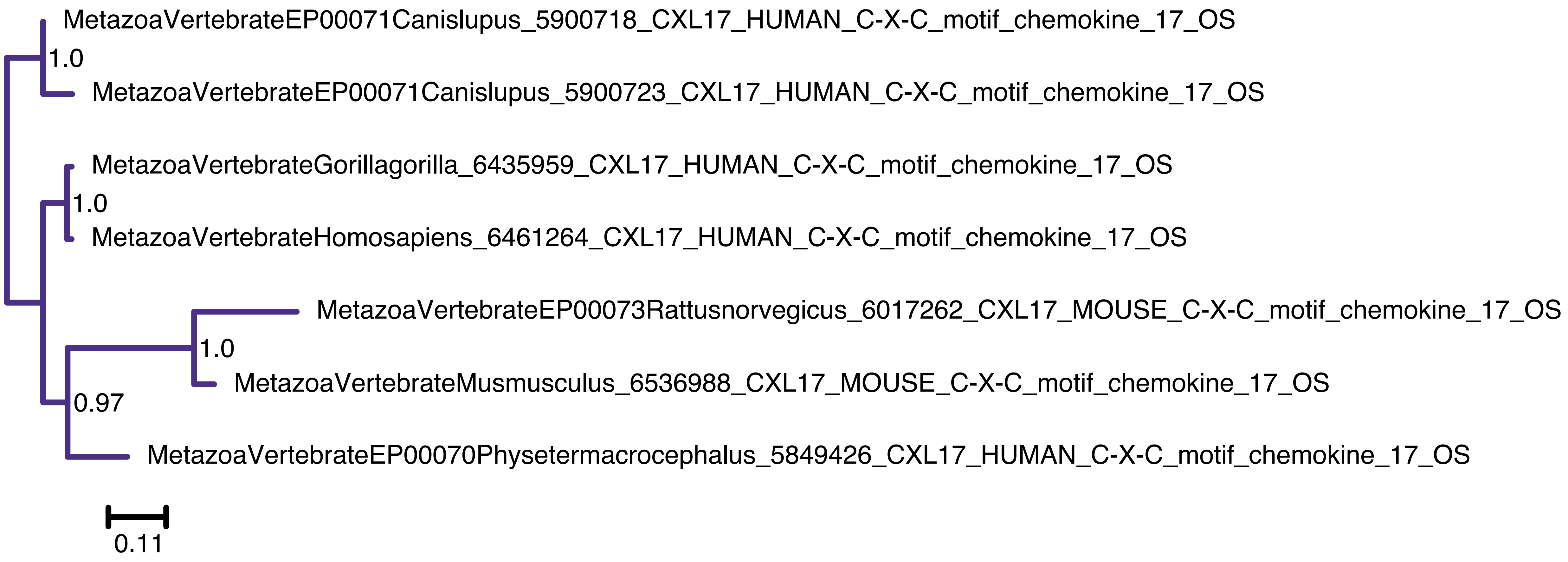

MetazoaVertebrateEP00071Canislupus_5900718_CXL17_HUMAN_C-X-C_motif_chemokine_17_OS
1.0
MetazoaVertebrateEP00071Canislupus_5900723_CXL17_HUMAN_C-X-C_motif_chemokine_17_OS
MetazoaVertebrateGorillagorilla_6435959_CXL17_HUMAN_C-X-C_motif_chemokine_17_OS
1.0
MetazoaVertebrateHomosapiens_6461264_CXL17_HUMAN_C-X-C_motif_chemokine_17_OS
MetazoaVertebrateEP00073Rattusnorvegicus_6017262_CXL17_MOUSE_C-X-C_motif_chemokine_17_OS
1.0
MetazoaVertebrateMusmusculus_6536988_CXL17_MOUSE_C-X-C_motif_chemokine_17_OS
0.97
MetazoaVertebrateEP00070Physetermacrocephalus_5849426_CXL17_HUMAN_C-X-C_motif_chemokine_17_OS
0.11

Fig. S15

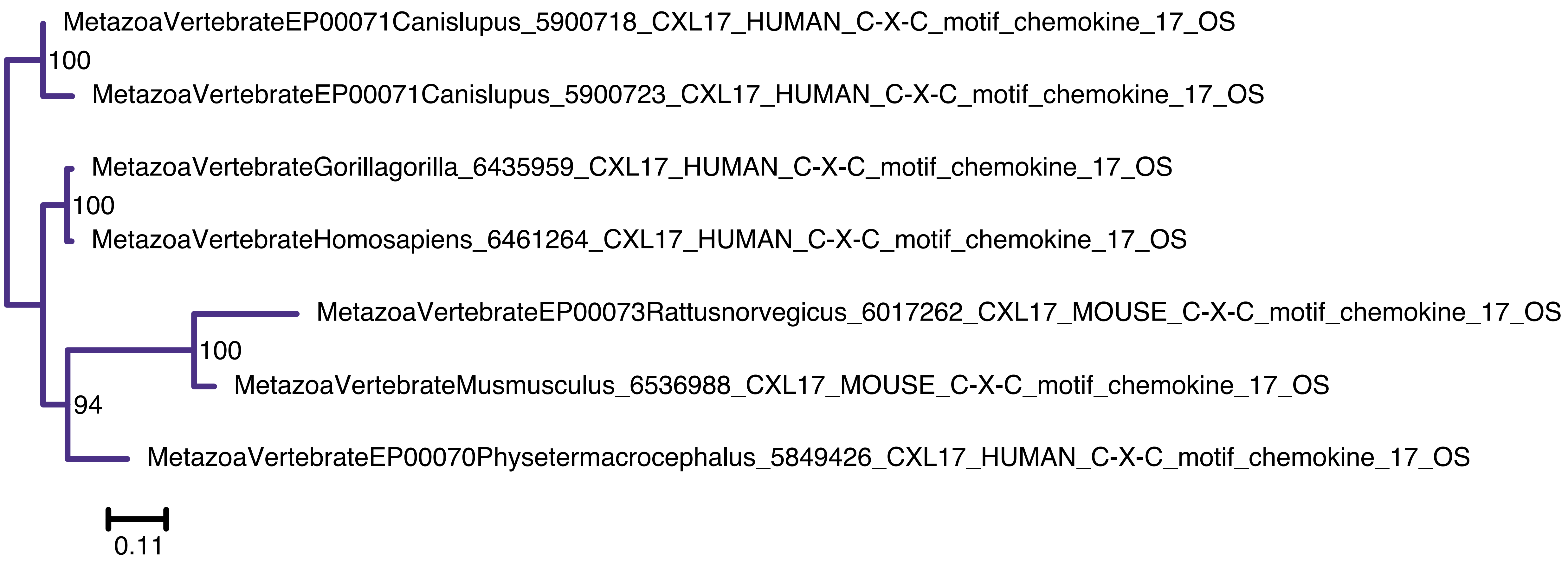
MetazoaVertebrateEP00071Canislupus_5900718_CXL17_HUMAN_C-X-C_motif_chemokine_17_OS
100
MetazoaVertebrateEP00071Canislupus_5900723_CXL17_HUMAN_C-X-C_motif_chemokine_17_OS
MetazoaVertebrateGorillagorilla_6435959_CXL17_HUMAN_C-X-C_motif_chemokine_17_OS
100
MetazoaVertebrateHomosapiens_6461264_CXL17_HUMAN_C-X-C_motif_chemokine_17_OS
MetazoaVertebrateEP00073Rattusnorvegicus_6017262_CXL17_MOUSE_C-X-C_motif_chemokine_17_OS
100
MetazoaVertebrateMusmusculus_6536988_CXL17_MOUSE_C-X-C_motif_chemokine_17_OS
94
MetazoaVertebrateEP00070Physetermacrocephalus_5849426_CXL17_HUMAN_C-X-C_motif_chemokine_17_OS
0.11

**Fig. S16**

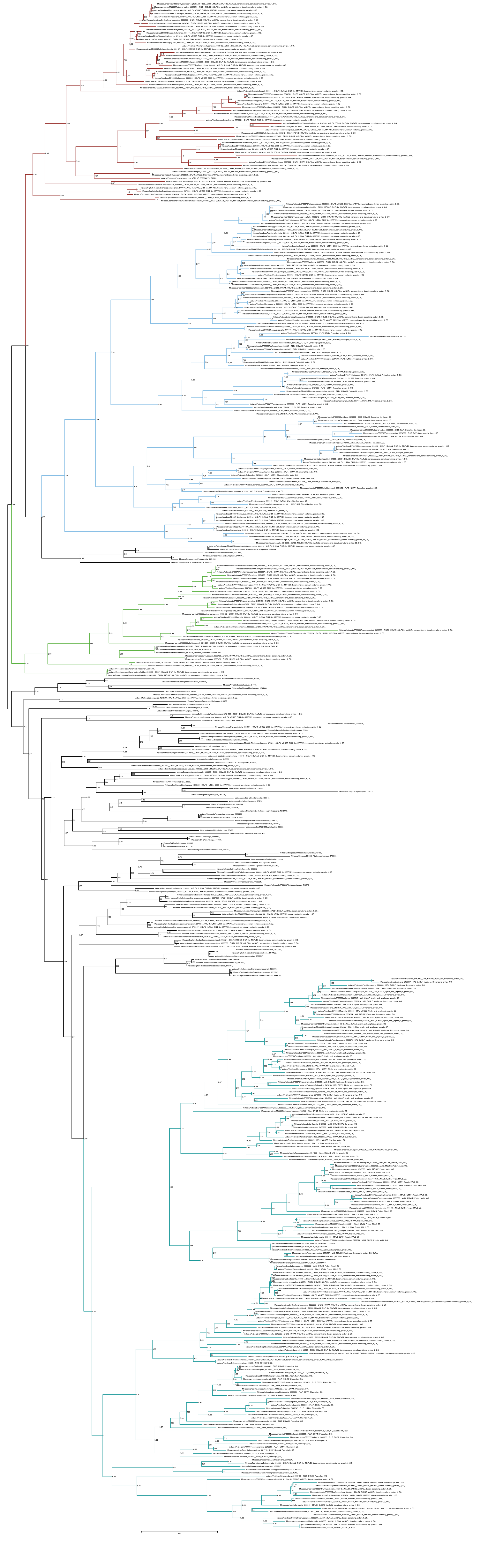

**Fig. S17**

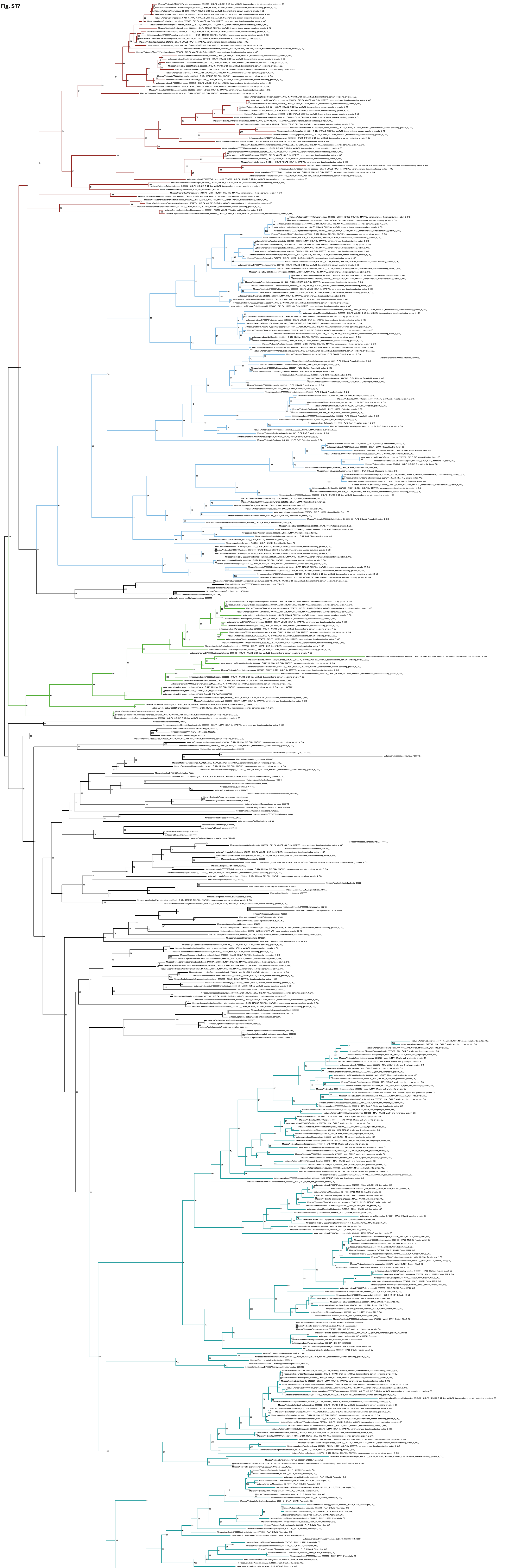

**Fig. S18**

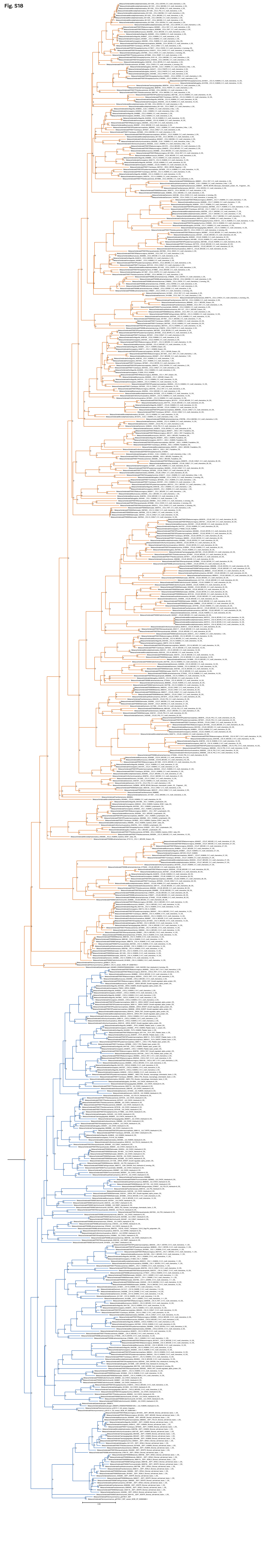

**Fig. S19**

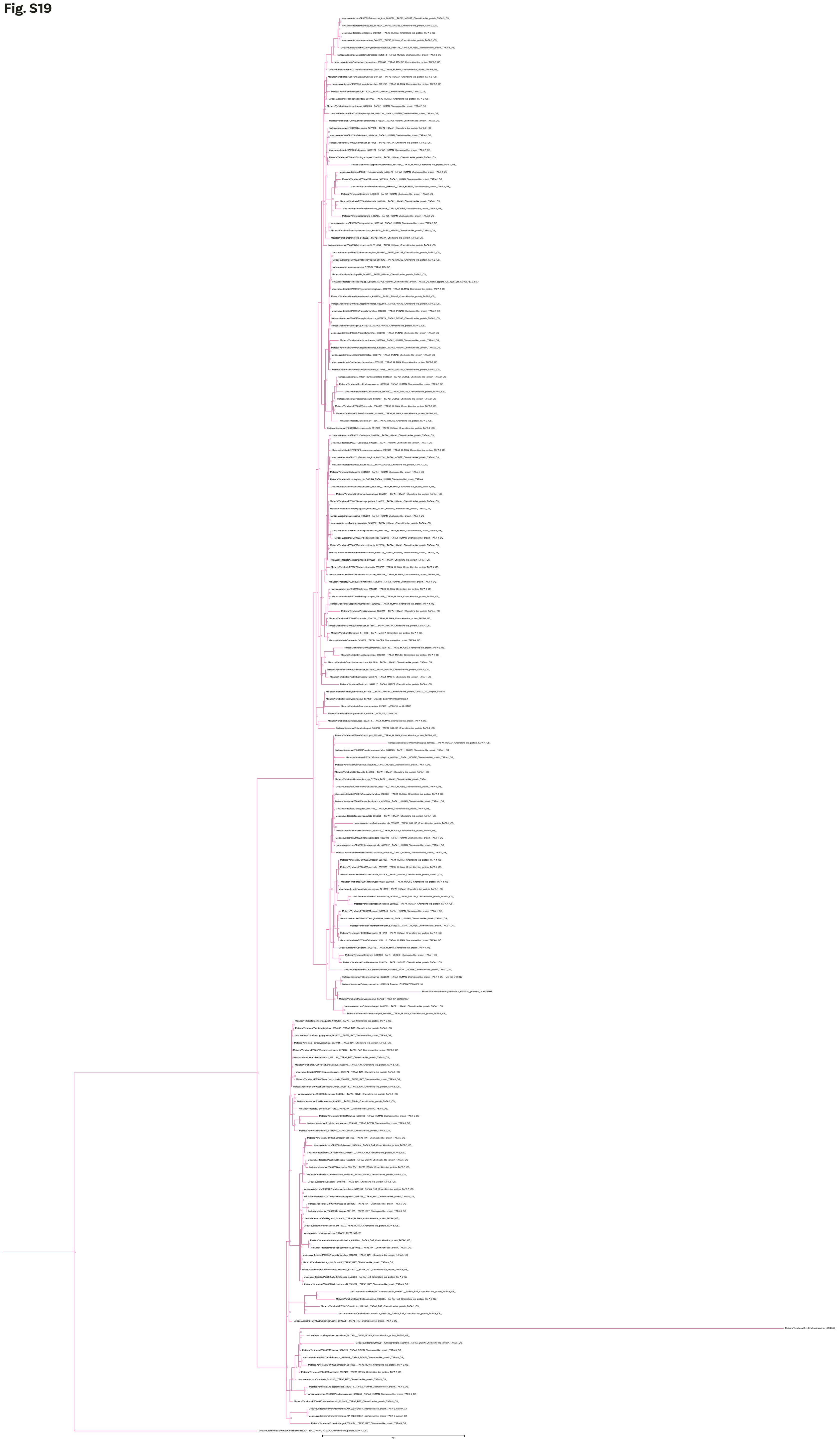

**Fig. S20**

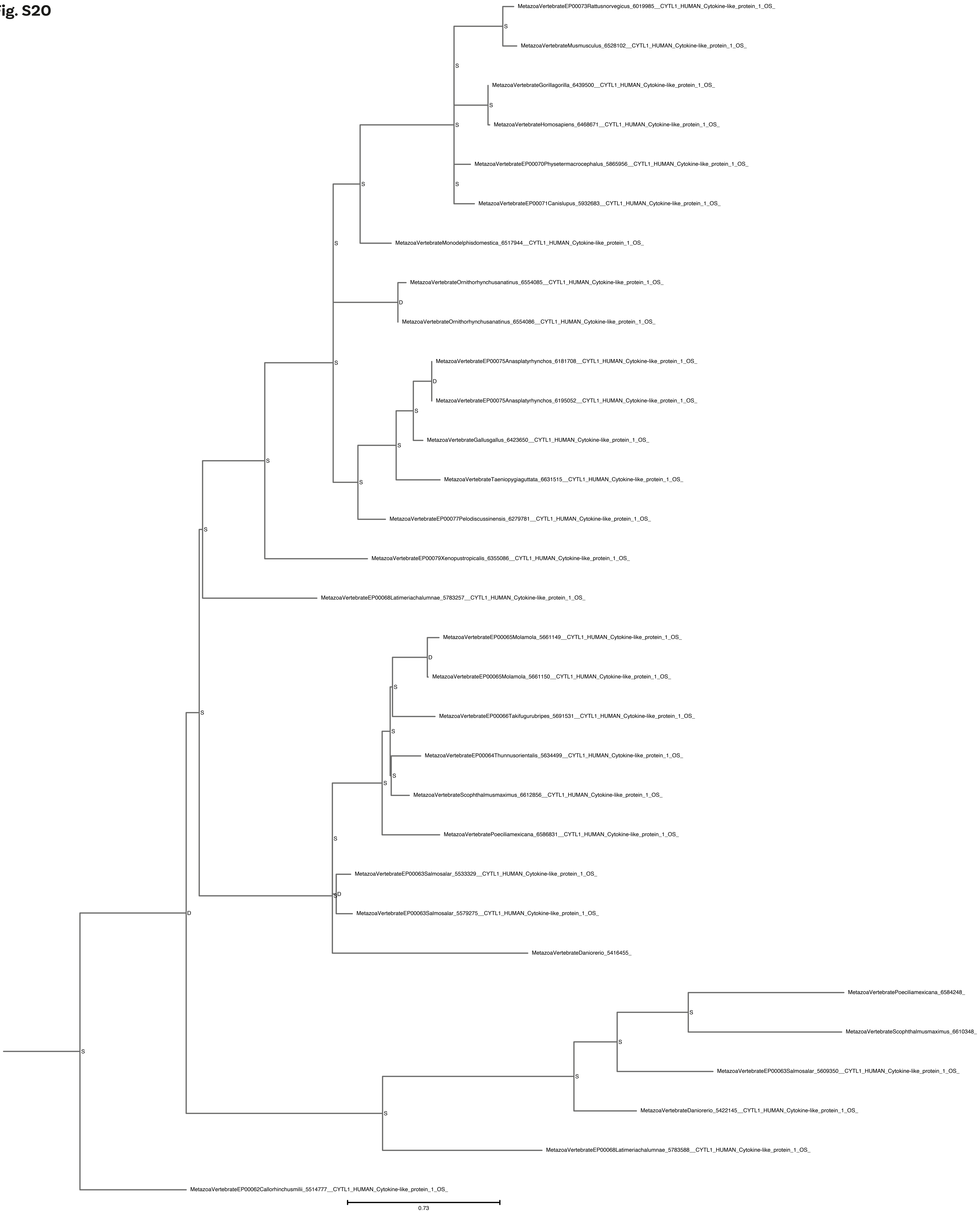
MetazoaVertebrateEP00073Rattusnorvegicus_6019985__CYTL1_HUMAN_Cytokine-like_protein_1_OS_
S
MetazoaVertebrateMusmusculus_6528102__CYTL1_HUMAN_Cytokine-like_protein_1_OS_
S
MetazoaVertebrateGorillagorilla_6439500__CYTL1_HUMAN_Cytokine-like_protein_1_OS_
S
S
MetazoaVertebrateHomosapiens_6468671__CYTL1_HUMAN_Cytokine-like_protein_1_OS_
MetazoaVertebrateEP00070Physetermacrocephalus_5865956__CYTL1_HUMAN_Cytokine-like_protein_1_OS_
S
S
MetazoaVertebrateEP00071Canislupus_5932683__CYTL1_HUMAN_Cytokine-like_protein_1_OS_
S
MetazoaVertebrateMonodelphisdomestica_6517944__CYTL1_HUMAN_Cytokine-like_protein_1_OS_
MetazoaVertebrateOrnithorhynchusanatinus_6554085__CYTL1_HUMAN_Cytokine-like_protein_1_OS_
D
MetazoaVertebrateOrnithorhynchusanatinus_6554086__CYTL1_HUMAN_Cytokine-like_protein_1_OS_
S
MetazoaVertebrateEP00075Anasplatyrhynchos_6181708__CYTL1_HUMAN_Cytokine-like_protein_1_OS_
D
MetazoaVertebrateEP00075Anasplatyrhynchos_6195052__CYTL1_HUMAN_Cytokine-like_protein_1_OS_
S
S
MetazoaVertebrateGallusgallus_6423650__CYTL1_HUMAN_Cytokine-like_protein_1_OS_
S
MetazoaVertebrateTaeniopygiaguttata_6631515__CYTL1_HUMAN_Cytokine-like_protein_1_OS_
S
MetazoaVertebrateEP00077Pelodiscussinensis_6279781__CYTL1_HUMAN_Cytokine-like_protein_1_OS_
S
MetazoaVertebrateEP00079Xenopustropicalis_6355086__CYTL1_HUMAN_Cytokine-like_protein_1_OS_
MetazoaVertebrateEP00068Latimeriachalumnae_5783257__CYTL1_HUMAN_Cytokine-like_protein_1_OS_
MetazoaVertebrateEP00065Molamola_5661149__CYTL1_HUMAN_Cytokine-like_protein_1_OS_
D
MetazoaVertebrateEP00065Molamola_5661150__CYTL1_HUMAN_Cytokine-like_protein_1_OS_
S
S
MetazoaVertebrateEP00066Takifugurubripes_5691531__CYTL1_HUMAN_Cytokine-like_protein_1_OS_
S
MetazoaVertebrateEP00064Thunnusorientalis_5634499__CYTL1_HUMAN_Cytokine-like_protein_1_OS_
S
S
MetazoaVertebrateScophthalmusmaximus_6612856__CYTL1_HUMAN_Cytokine-like_protein_1_OS_
MetazoaVertebratePoeciliamexicana_6586831__CYTL1_HUMAN_Cytokine-like_protein_1_OS_
S
MetazoaVertebrateEP00063Salmosalar_5533329__CYTL1_HUMAN_Cytokine-like_protein_1_OS_
D
S
MetazoaVertebrateEP00063Salmosalar_5579275__CYTL1_HUMAN_Cytokine-like_protein_1_OS_
D
MetazoaVertebrateDaniorerio_5416455_
MetazoaVertebratePoeciliamexicana_6584248_
S
MetazoaVertebrateScophthalmusmaximus_6610348_
S
S
MetazoaVertebrateEP00063Salmosalar_5609350__CYTL1_HUMAN_Cytokine-like_protein_1_OS_
S
MetazoaVertebrateDaniorerio_5422145__CYTL1_HUMAN_Cytokine-like_protein_1_OS_
S
MetazoaVertebrateEP00068Latimeriachalumnae_5783588__CYTL1_HUMAN_Cytokine-like_protein_1_OS_
MetazoaVertebrateEP00062Callorhinchusmilii_5514777__CYTL1_HUMAN_Cytokine-like_protein_1_OS_
0.73

**Fig. S21**

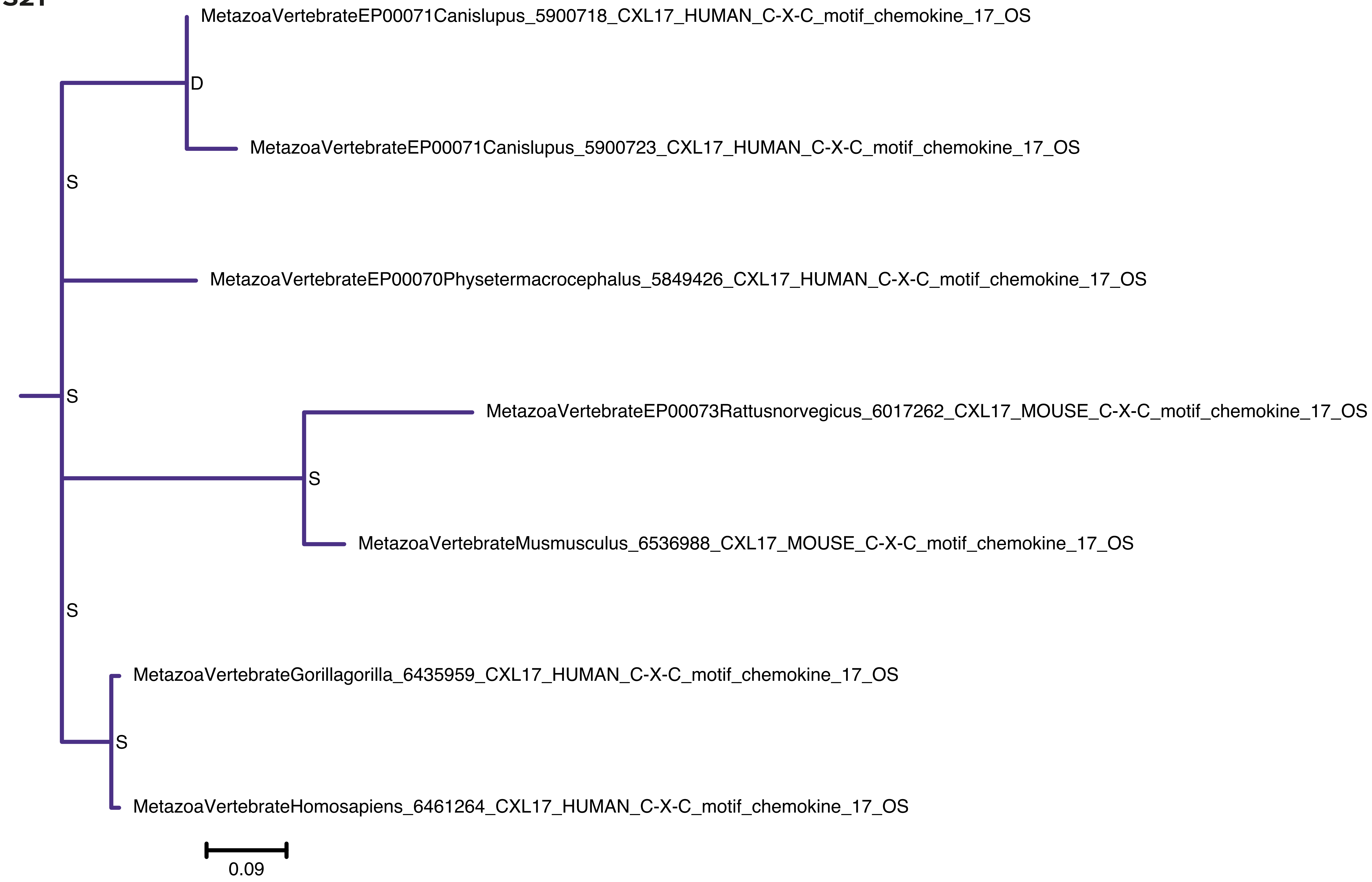
MetazoaVertebrateEP00071Canislupus_5900718_CXL17_HUMAN_C-X-C_motif_chemokine_17_OS
D
MetazoaVertebrateEP00071Canislupus_5900723_CXL17_HUMAN_C-X-C_motif_chemokine_17_OS
S
MetazoaVertebrateEP00070Physetermacrocephalus_5849426_CXL17_HUMAN_C-X-C_motif_chemokine_17_OS
S
MetazoaVertebrateEP00073Rattusnorvegicus_6017262_CXL17_MOUSE_C-X-C_motif_chemokine_17_OS
S
MetazoaVertebrateMusmusculus_6536988_CXL17_MOUSE_C-X-C_motif_chemokine_17_OS
S
MetazoaVertebrateGorillagorilla_6435959_CXL17_HUMAN_C-X-C_motif_chemokine_17_OS
S
MetazoaVertebrateHomosapiens_6461264_CXL17_HUMAN_C-X-C_motif_chemokine_17_OS
0.09

### Fig. S22

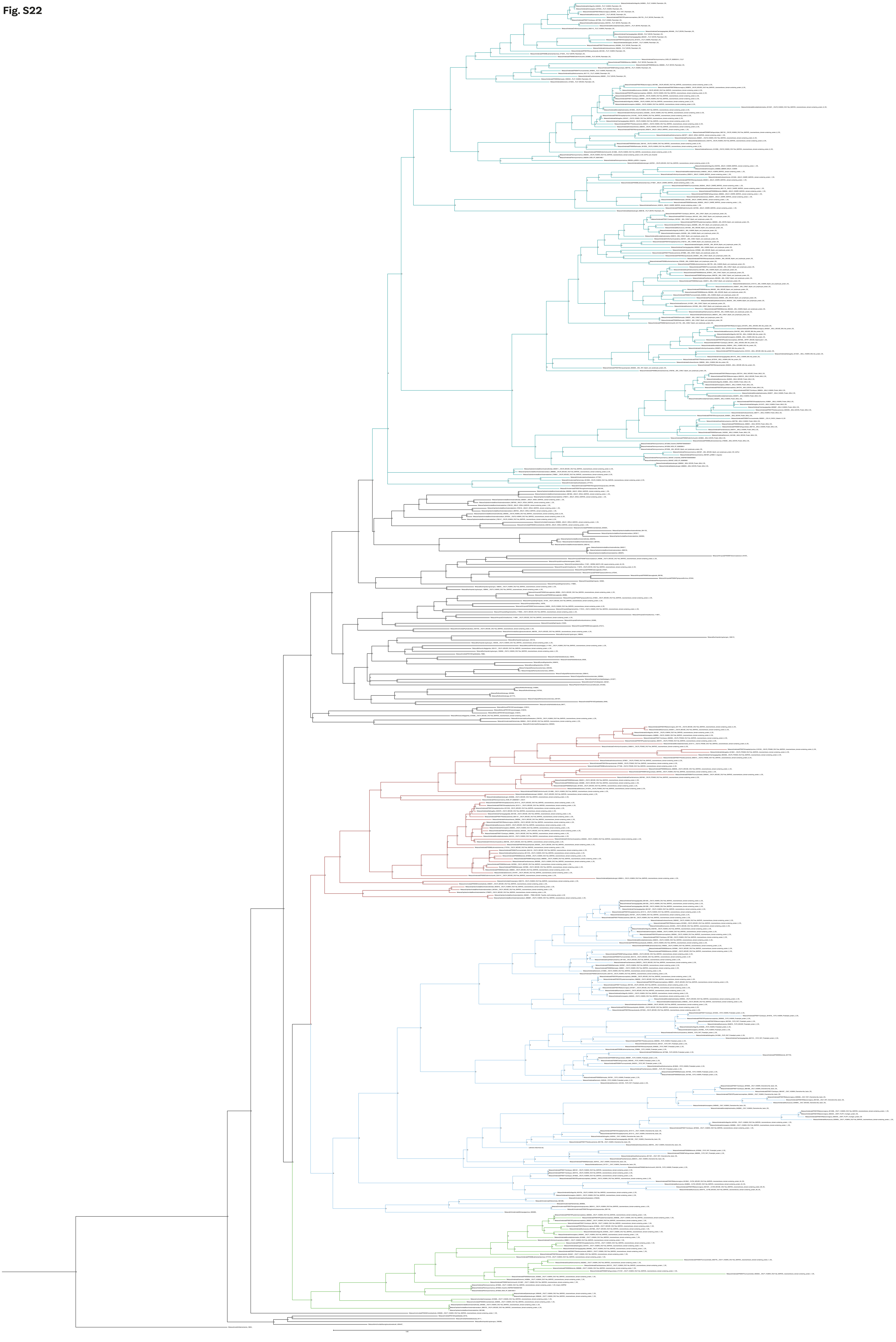

#### Fig. S23

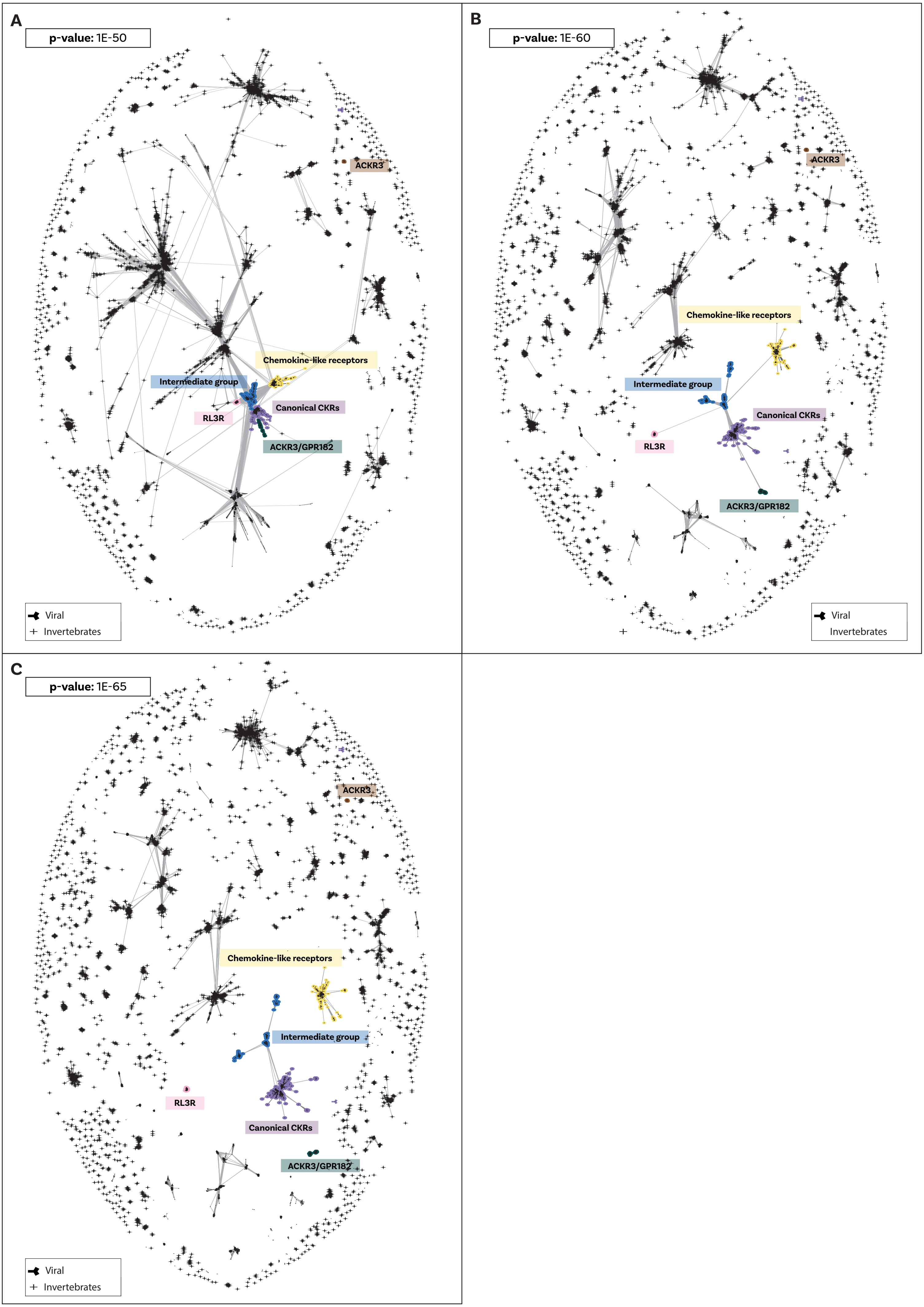

A
p-value: 1E-50
Chemokine-like receptors
Intermediate group
Canonical CKRs
RL3R
ACKR3/GPR182
ACKR3
Viral
Invertebrates
B
p-value: 1E-60
ACKR3
Chemokine-like receptors
Intermediate group
Canonical CKRs
RL3R
ACKR3/GPR182
Viral
Invertebrates
C
p-value: 1E-65
ACKR3
Chemokine-like receptors
Intermediate group
RL3R
Canonical CKRs
ACKR3/GPR182
Viral
Invertebrates

**Fig S24**

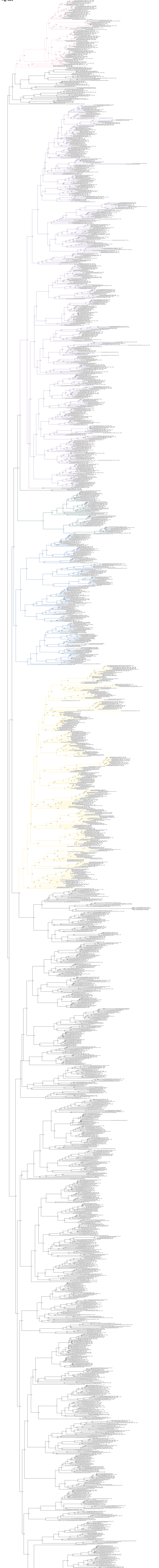

**Fig S25**

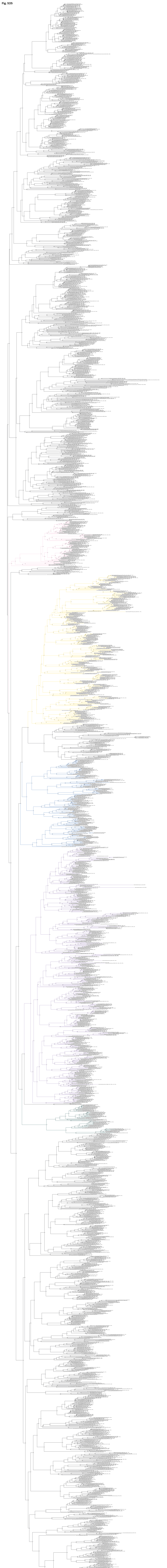

**Fig. S26**

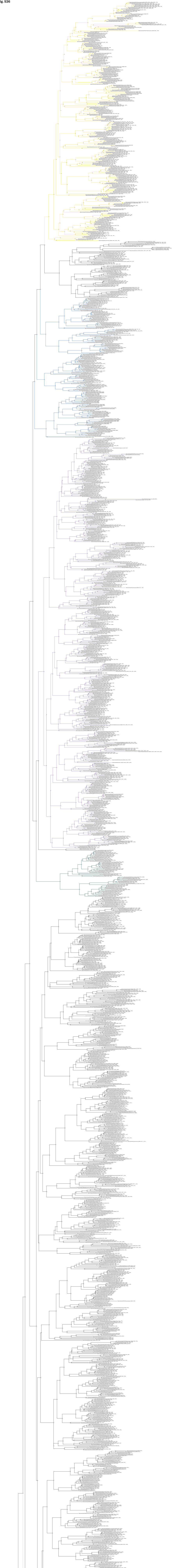
