## Supplementary results for "The origin, evolution and molecular diversity of the chemokine system"

**Exploration of candidate invertebrate homologs to canonical chemokines and related molecules**

To explore the possibility of finding canonical chemokines and/or related molecules (TAFA and CYTL) outside of vertebrates, we used BLASTP with a loose e-value threshold of 0.1 to search 39 invertebrate proteomes (see Table S2). 18 candidate sequences were collected and explored further.

From the clustering analysis in CLANS (Figure 1A, Figure S1 and File S1), it became apparent that 4 sequences were candidate TAFAs, while the remaining 14 sequences connected loosely to the canonical chemokines. We then performed BLASTP (1, 2) versus the curated SwissProt dataset (3, 4) and collected the first 5 hits. Furthermore, we used InterProScan (5, 6) to identify protein signatures. See Supplementary File S3 for a summary of all these results for all sequences.

Regarding the canonical chemokines, only 3 sequences received annotations related to chemokines from the BLASTP versus SwissProt. These were: one brachiopod sequence (*Lingula unguis*) as candidate CCL24, one cnidarian (*Clytia hemisphaerica*) sequence as candidate CCL3, and one echinoderm (*Acanthaster planci*) sequence as candidate CXCL10. Although none of these sequences were categorised as chemokines with InterProScan, we anyway decided to look at them further. Firstly, we noted how all three sequences were significantly longer than their counterparts in vertebrates. Secondly, none of the three sequences possessed a signal peptide, as calculated with SignalP 6.0 online tool (<https://services.healthtech.dtu.dk/service.php?SignalP-6.0>), which is expected from secreted proteins such as chemokines (7). Finally, we anyway tried to align the sequences (MAFFT –auto) with their respective candidate relatives and found poor conservation (Figures S3-5). The lack of evidence for being true chemokines, led us to discard all invertebrate candidates for further analyses on canonical chemokines.

The 4 candidate invertebrate TAFA sequences all belong to the urochordate *Ciona intestinalis*. One sequence was annotated as TAFA by both SwissProt and InterProScan, while the other three appear to be prolyl hydroxylases (see Supplementary File S3). We anyway studied all four sequences further and found that the only one to possess a signal peptide is the sequence that received TAFA annotation. Moreover, the other three sequences appear to be too long and poorly aligned with vertebrate TAFAs (Figure S6). The TAFA annotated sequence was of correct length and showed sequence conservation in the alignments (Figures S6-S7). Interestingly, it possesses 8 of the 10 typical cysteine residues of TAFA1-4 and the two missing cysteines are the same ones missing in TAFA5. Considering that TAFA5 is sister group to TAFA1-4 and that the urochordate sequence places itself as orthologous to all TAFAs (see main text Results) it is reasonable to conclude that the ancestral TAFAs possessed 8 cysteine residues and that the additional cysteines are a novelty of the TAFA1-4 lineage.

Taken together these results show that while canonical chemokines are indeed a vertebrate innovation, TAFA “chemokines” likely originated in the ancestor of vertebrates and urochordates.

**Exclusion of some sequences from the CKLFSF dataset**

The data mining through BLASTP and PSI-BLAST (8) provided numerous candidate CKLFSF homologs both in vertebrates and in invertebrates and these were all included in a clustering analysis with CLANS (Figure 1C and Figure S2). Two main clusters emerged and we called them “CKLF I” and “CKLF II”. While CKLF I was vertebrate specific, the CKLF II included multiple invertebrate sequences. While the two main clusters are well-defined already at p-values of ~1E-20 (Figure S2 and File S2), they only connect to each other at 1E-15. At this p-value, 4 additional sequences connected loosely to the CKLF II cluster. These were 3 echinoderm sequences (all from *Stichopus japonicus*) and one sequence from the placozoan *Trichoplax adhaerens*. The latter is the only non-bilaterian sequence collected from the original BLASTs. These sequences not only joined the CKLFSF cluster at the limit threshold, but also connected only to sequences that were already marginal, therefore being only indirectly connected to the core of the cluster. Like above, we examined the sequences with a BLAST versus SwissProt and with InterProScan (see Supplementary File S3). The evidence in favour of keeping these sequences was scant (see details in Supplementary File S3), and we decided to exclude them from downstream phylogenetic analyses. The CKLFSF dataset therefore did not include any non-bilaterian sequences, although multiple bilaterian invertebrate phyla were represented.
